# Spike-wave discharges reflect widespread, but non-uniform, cortical hypersynchrony during immobility

**DOI:** 10.64898/2026.09.14.751500

**Authors:** Scott Kilianski, Elzbieta Dulko, Anna Grace Carns, Sakina Lashkeri, Magdalena Pikus, Mark Beenhakker

**Author notes:** Neuroscience Graduate Program, Baylor College Medicine, Baylor College of Medicine, 1 Baylor Plaza, Houston, TX 77030, USA. Department of Neuroscience, Karolinska Institutet, C4 Neurovetenskap, C4 Forskning Stagkourakis, 171 77 Stockholm, Sweden (Magdalena Pikus).

## Abstract

Spike-wave discharges (SWDs) are the electrographic hallmark of absence epilepsy, yet direct evidence for their proposed basis in cortical hypersynchrony has been scarce. We addressed this gap by recording neuronal populations across three cortical regions — primary somatosensory (S1), visual (V1), and secondary motor cortex (M2) — using silicon probe arrays in spontaneously seizing C3H/HeJ mice. SWDs drove profound increases in neuronal synchrony, rhythmicity, and phase-locking across all recorded regions. V1 and M2 neurons were clearly entrained to SWD cycles, though less strongly than S1 neurons. Electrical stimulation of both S1 and V1 could induce or terminate SWDs, supporting widespread network involvement. Finally, we also provide evidence that SWD-associated reductions in firing rate could be attributable to co-occurring behavioral immobility rather than to the seizures themselves.

**Graphical Abstract:** 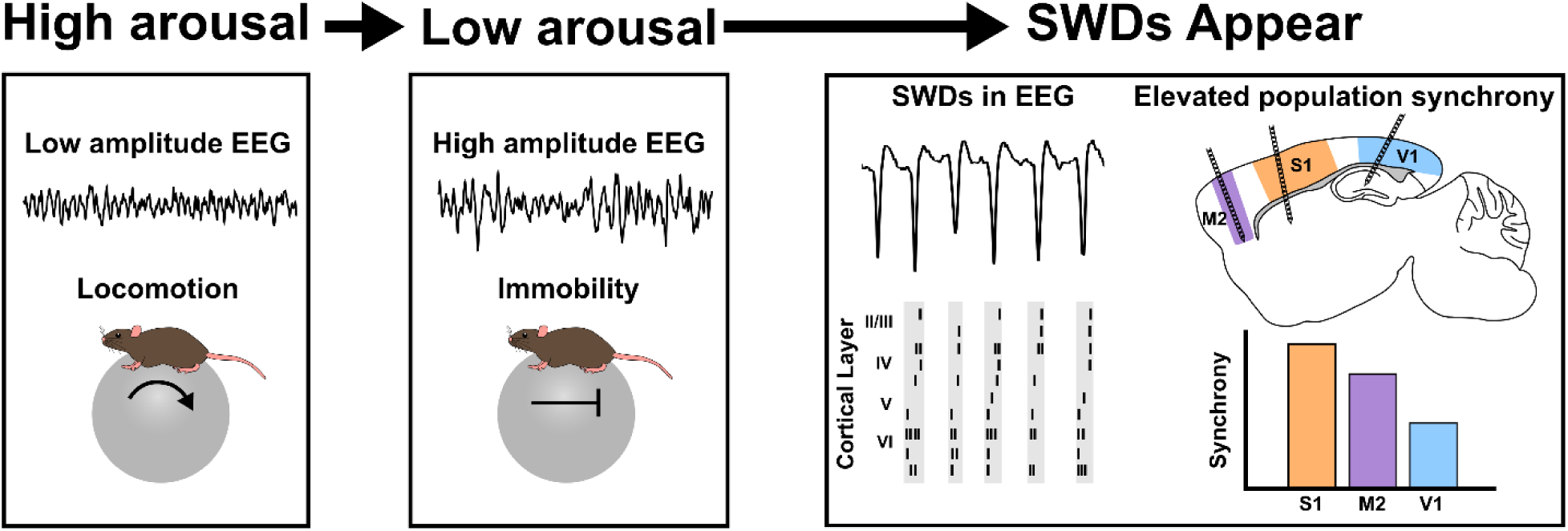

## INTRODUCTION

Absence seizures are sudden, brief lapses in awareness accompanied by high amplitude, periodic spike-wave discharges (SWDs) recorded in the electroencephalogram (EEG) (Blumenfeld, 2005a, 2005a; Coenen, 2003; Gibbs et al., 1935; Panayiotopoulos, 2008). Despite diverse genetic etiologies (Balestrini et al., 2026; Crunelli & Leresche, 2002), SWDs are a universal feature of absence epilepsy, and thus represent a stereotyped signature of cortical dynamics associated with absence seizures. Although SWDs are commonly attributed to hypersynchronous cortical spiking, direct evidence that neuronal population synchrony *per se* is elevated during SWDs is surprisingly sparse.

While many studies in rodent models of absence epilepsy have recorded the activity of single neurons (Khan et al., 2024; McCafferty et al., 2018, 2023; Paz et al., 2005, 2007; Pinault, 2003; Pinault et al., 1998; Polack et al., 2007; Polack & Charpier, 2006) or aggregate multi-unit signals (Buzsaki et al., 1988; Inoue et al., 1993; Kandel & Buzsáki, 1997; Terlau et al., 2020), such recordings cannot resolve population-level synchrony (Salinas & Sejnowski, 2001). Although large-scale neuronal recordings during SWDs are emerging in the literature (Kozák et al., 2020; McCafferty et al., 2018, 2023; Meyer et al., 2018), studies that have directly assessed synchrony have, surprisingly, found minimal (motor/prefrontal cortex: Kozák et al., 2020) or even reduced (visual cortex: Meyer et al., 2018) synchrony during SWDs relative to baseline. These discrepant results may reflect differences in the cortical regions examined or the animal models used (i.e., rat versus mouse). Moreover, quantitative assessments of neuronal synchrony in somatosensory cortex, the presumed site of SWD initiation, are lacking. We therefore sought to systematically quantify neuronal synchrony across multiple cortices within a single rodent model of absence seizures.

To resolve the magnitude and ubiquity of cortical activity and synchrony during SWDs, we recorded neuronal populations in three cortical areas spanning most of the anteroposterior axis: primary somatosensory (S1), primary visual (V1), and secondary motor cortex (M2) in C3H/HeJ mice. This model of absence epilepsy has frequent, spontaneous SWDs due to a mutation in the *Gria4* gene encoding the GluR4 AMPA receptor subunit (Beyer et al., 2008; Frankel et al., 2005). We first characterized the recruitment of these different cortical regions to SWDs by measuring neuronal firing rates, neuronal phase-locking, consistency of activated neuronal sequences, and population-level synchrony during SWDs. We also directly electrically stimulated S1 and V1 to determine if either region could induce or interrupt SWDs. Our findings demonstrate that SWDs recruit widespread synchrony across cortical regions, including V1, contradicting earlier findings (Meyer et al., 2018). We also found that electrically stimulating either S1 or V1 can control SWDs. Finally, we report that SWDs are associated with specific patterns of cortical laminar activation that preferentially occur during behavioral arrest.

## RESULTS

### Recording neural activity in head-fixed C3H/HeJ mice having spontaneous SWDs

To evaluate the relationship between spontaneous SWDs and neuronal activity patterns, we performed acute, silicon probe recordings in awake, head-fixed C3H/HeJ mice (**Figure 1A**). Mice were implanted with a surface EEG electrode on the cortical surface under the frontal skull bone and outfitted with headbars for subsequent head-fixation on a rotatable ball. To measure the activity of single cortical neurons (i.e., *cortical units*), we performed silicon probe recordings (Shobe et al., 2015) that targeted either S1, V1, or M2 cortex (**Figure 1B**). Altogether, we obtained 37 one-hour recording sessions from 24 mice; in a subset of mice, activity was recorded in two separate sessions. Cortical units and surface EEG were simultaneously recorded (**Figure 1C**) and histology was performed to verify the location of the cortical units (**Figure 1D**). Regardless of cortical target, the surface EEG electrode under the frontal bone was used to define SWD start and end times, as well as the timepoints of individual SWD spikes (i.e., prominent troughs during the seizures). SWD rates, durations, and peak frequencies were indistinguishable across recording sessions (one-way ANOVA *p* > 0.05, **Figure 1E**, **Table S1**).

**Figure 1.**
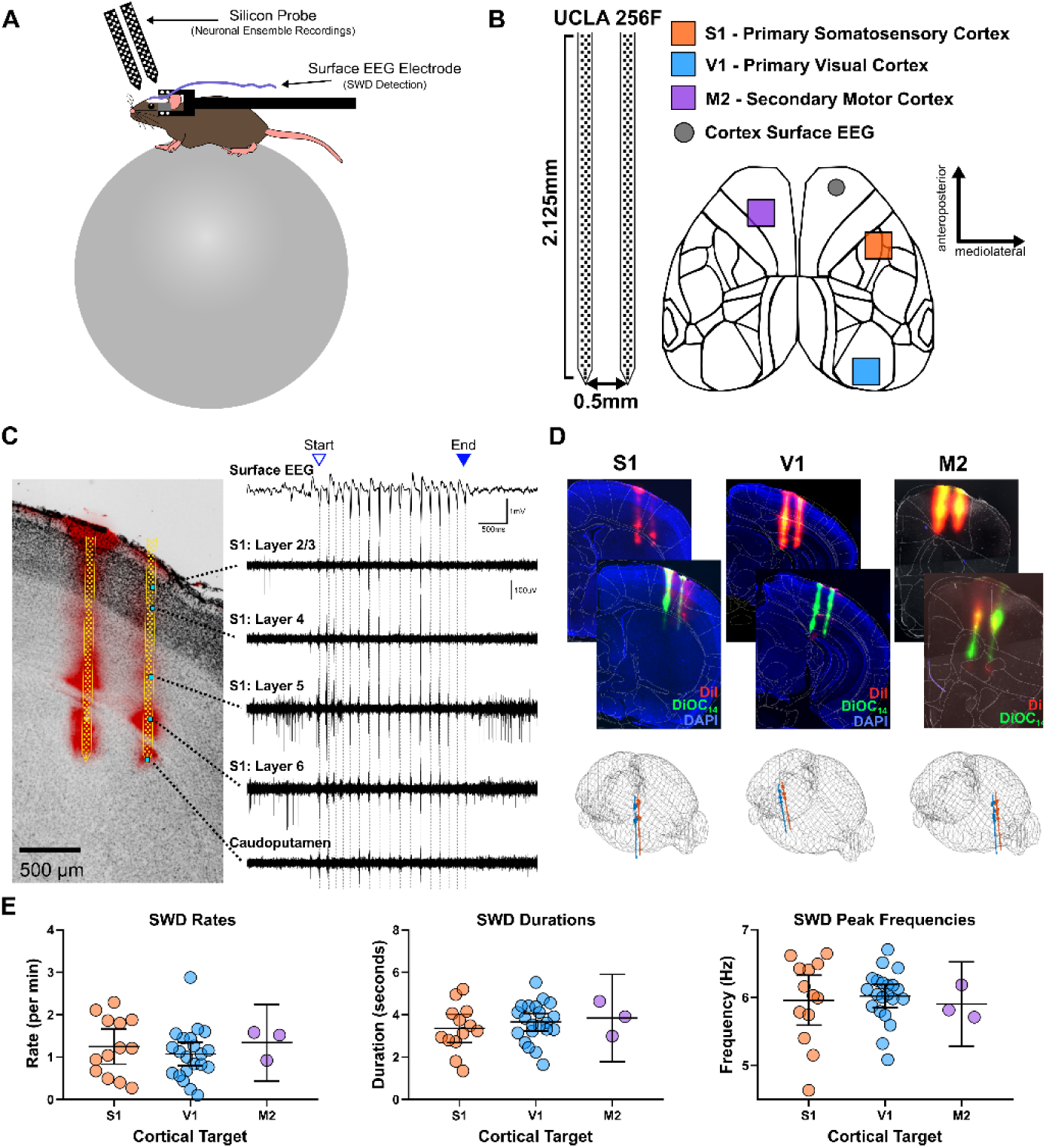
Head-fixed silicon probe recordings. **(A)** Schematic of head-fixed recording apparatus for acute, awake silicon probe recordings. A mouse is affixed to a rod allowing for voluntary locomotion while keeping the head in place. EEG is used to detect SWDs. **(B)** Left: UCLA 256F silicon probe diagram showing the vertical electrode span (2.125mm) on two shanks separated by 0.5mm. Right: Top-down view of the mouse cortex. Sites for silicon probe insertions are shown in color and surface EEG in gray. **(C)** Left: Example histology with scaled probe map overlaid in yellow. Right: Surface EEG and high-pass (>300Hz) filtered traces from corresponding silicon probe channels labeled in cyan (Left) during SWD. Triangles mark the start and end of the SWD. Dashed gray lines mark individual SWD troughs. **(D)** Top: Representative histology from recordings in S1, V1, and M2 cortex using two dyes with distinct emission/excitation profiles: DiI (red) and DiOC14 (green). Bottom: 3D wireframe brain models displaying the best-fit line through the brain based on the probe path reconstruction procedure. Circles show user-drawn labels used in the procedure. **(E)** Average SWD rates, durations, and peak frequencies in recordings at each cortical target site. Each circle is the average value for a single recording session. There was no significant effect of cortical target on any SWD feature (one-way repeated measures ANOVAs, all p > 0.05). Error bars indicate 95% confidence intervals.

### Behavioral arrest largely accounts for SWD-associated firing rate changes

Recent work in the GAERs rat model of absence epilepsy has surprisingly demonstrated that firing rates among the majority of neurons in the somatosensory cortex are either modestly suppressed or unchanged several seconds prior to SWD onset, and that such suppression generally persists during the SWD (McCafferty et al., 2023). Similarly, average firing rates among somatosensory cortex neurons in the C3H/HeJ mice recorded during this study appeared to only decrease modestly during SWDs (**Figure 2A**). This finding was preserved when the 0.5s bin duration employed in previous studies was applied (**Figure S1**). We found little apparent firing rate change among V1 or M2 neurons during SWDs (**Figure 2A**). Ultimately, across all cortical regions, firing rates during SWDs were subtly, but significantly, lower when compared to rates during all non-ictal, baseline periods (**Figure S1B**). Firing rates decreased in S1 (-0.86 spikes/s, n = 785 neurons, *p* = 5.0×10^−9^, paired t-test), M2 (-0.26 spikes/s, n = 350 *p* = 0.018), and V1 (-0.64 spikes/s, n = 513, *p* = 1.0×10^−6^). As our electrodes also captured activity ventral to the cortex, we learned that firing rate reductions are not universally observed: such rates increased in the caudoputamen (**Figure S1**).

**Figure 2.**
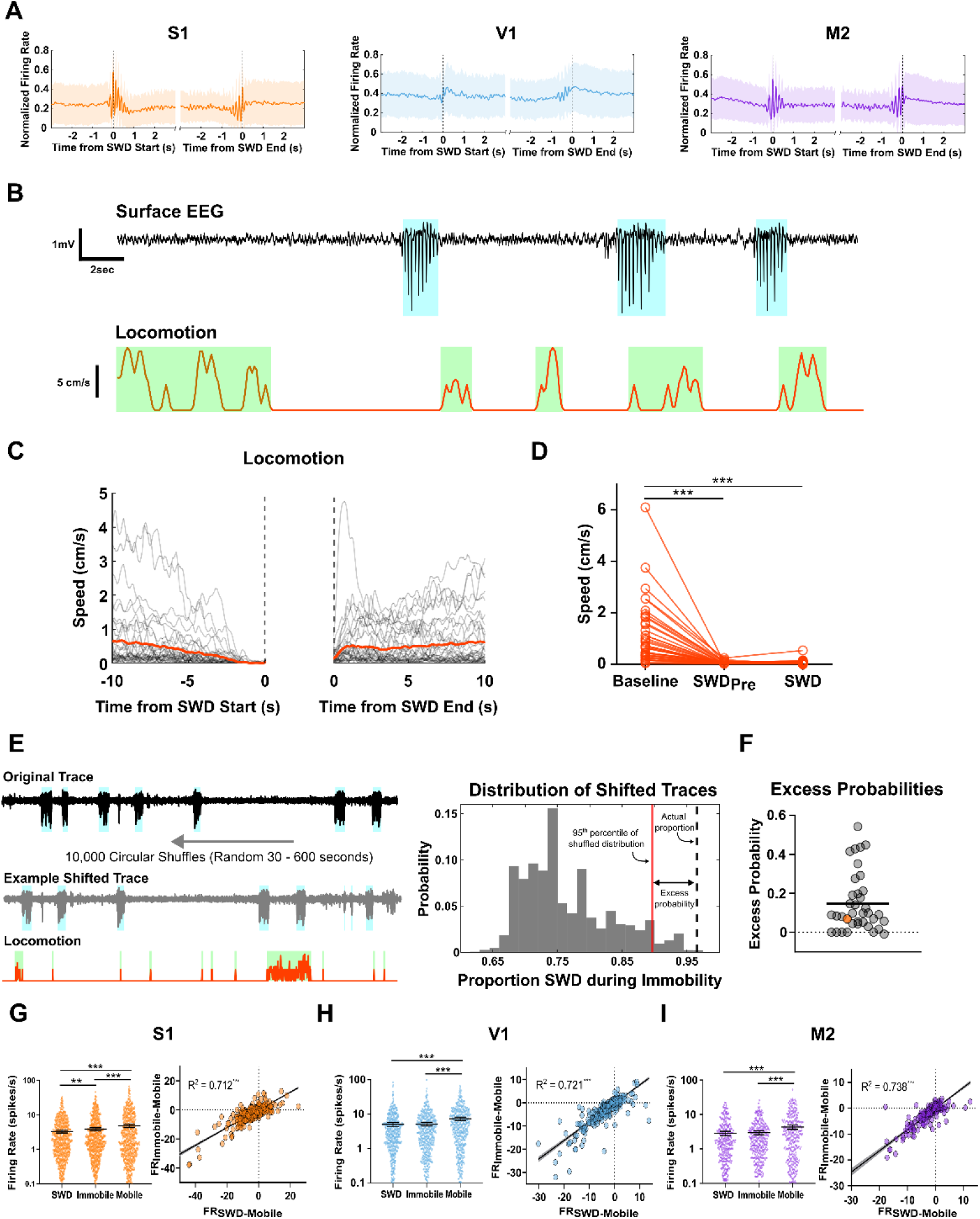
Firing rates subtly decrease during SWDs and Immobility. **(A)** Average normalized firing rates relative to the starts (left) and ends (right) of SWDs. Shading is ± SD. **(B)** Example traces of EEG and locomotion. Cyan and green shading indicates SWD and movement, respectively. **(C)** Left: Locomotion (speed) relative to starts and ends of SWDs. Grey lines show mean traces from individual sessions. Thick red lines show mean of all sessions. **(D)** Average movement speed during baseline, SWDPre, and SWD periods. **(E)** Left: Example of temporal shifting procedure. EEG traces were circularly temporally shifted 30 to 600 seconds while locomotion traces were unaltered. The proportion of SWDs starting during immobility was then determined. Right: The resultant distribution of the shuffling procedure (estimated null distribution) in gray bars. Red line indicates the value of the 95th percentile of the null distribution while the dotted black line corresponds to the observed proportion in the original data. The excess probabilities shown in Panel F correspond to the difference between the 95th percentile value and the observed proportion. **(F)** Excess probabilities for each session are shown as circles (n=37 sessions, 24 mice). The horizontal line indicates the mean across all sessions. The orange circle represents the difference score for the example session plotted in F. **(G-I)** Left: Firing rates of neurons during SWD, Immobile, and Mobile epochs for S1 (**G**, n = 785 neurons), V1 (**H**, n = 513 neurons), and M2 (**I**, n = 350 neurons). Horizontal black lines and bars show mean and 95% CI. Right: Scatter plots of firing rate differences between SWD/Immobile and Moving epochs. The positive slope and large R^2^ values indicate that firing rate changes observed during SWDs are similar to those observed during Immobility. Grey shading is 95% CI. ** p < 0.01, *** p < 10^−3^. S1 (n=386). V1 (n=358). M2 (n=244).

Although cortical firing rates decreased during SWDs, relative to baseline periods, we hypothesized that such reductions are epiphenomenal and largely reflect the behavioral arrest that accompanies SWDs; importantly, neuronal firing rates are modulated by motion (Harris & Thiele, 2011; McGinley et al., 2015; Polack et al., 2013; Saleem et al., 2013). Indeed, the co-occurrence of behavioral arrest and absence seizures is well established in both humans and experimental models (Blumenfeld, 2005b), and our recording sessions confirmed that SWDs primarily occur during periods of immobility (**Figure 2B**). Mouse locomotion gradually slowed in the seconds preceding SWD onset, culminating in complete immobility at seizure onset (**Figure 2C**, left). Mouse locomotion resumed after SWD termination **(Figure 2C**, right).

We compared mouse locomotion speed during different experimental epochs: (1) during SWDs (SWD), (2) 2.5s preceding SWDs (SWD_Pre_), and (3) all other times not contaminated with SWDs (Baseline). Indeed, one-way repeated measures ANOVA revealed that there was a significant effect of experimental epoch (i.e. Baseline, SWD_Pre_, SWD) on locomotion (F = 30.8, *p* = 2.2×10^− 10^). Locomotion was significantly reduced during SWD_Pre_ and SWD relative to baseline (*p* < 10^−7^, Bonferroni-corrected *post hoc* tests, **Figure 2D**) but not significantly different from each other (*p* > 0.99). Although these results indicate that a reduction in mouse motion occurs just prior to SWD generation, these results do not conclusively demonstrate that mice are already completely immobile when SWDs begin. We therefore analyzed the observed probability of SWDs occurring during immobility. Using the sensitive movement threshold of 1cm/s (see *Calculation of firing rates in different behavioral states* in STAR Methods), we found that > 98% (2788/2824) of all SWDs initiated while mice were already immobile. To test the likelihood of achieving the observed probabilities by chance in each recording, we applied a random temporal shifting procedure and computed an “excess probability” metric (**Figure 2E**). Excess probabilities were nearly always higher than the chance threshold (36/37 sessions), indicating that SWDs occur during immobility far more than expected by chance. Indeed, we speculate that the few SWDs detected during locomotion (36/2824) were caused by reflexive postural adjustments that spuriously triggered the rotary encoder and not by *bona fide* bouts of locomotion. In conclusion, SWDs occur nearly exclusively during complete immobility, consistent with the prevailing observation that behavioral arrest accompanies SWDs (Dong et al., 2024).

As SWDs coincide with behavioral arrest, the firing rate changes observed during SWDs may result from immobility and not from SWDs *per se*. To test this possibility, we partitioned recording sessions into three behavioral states: *SWD*, *Immobile*, and *Mobile*; the *Immobile* state did not include any behavioral arrests associated with SWDs. We then compared firing rates across these three states. In all recorded cortices (i.e., S1, V1, M2), firing rates decreased during both the *SWD* and *Immobile* states, relative to the *Mobile* state (*p* < 10^−4^ for all structures, Bonferroni’s multiple comparisons tests, **Figure 2G,H,I**). However, only S1 exhibited a significant difference in firing rate between the SWD and Immobile states. We also repeated these comparisons using the mean firing rate of all neurons in a recording session, rather than individual neurons, and found similar results (**Figure S1C**).

To further isolate the contribution of immobility to firing rate reductions, we calculated firing rate differences between states and performed linear regression. Firing rates during periods of immobility were subtracted from those of mobility to compute the firing rate difference between those states (FR_Immobile-Mobile_). We found that between 57% and 88% of the variance in SWD-related firing rate changes is shared by immobility (*R^2^* = 0.57 to 0.88, β = 0.60 to 0.84, p <10^−4^ for all structures, **Figure 2G,H,I**). As our silicon probes extended into subcortical regions, we similarly quantified the firing rate-behavioral state relationship in caudate putamen, subiculum, and the CA1 region of hippocampus, where we also found that neuronal firing rate changes largely reflect changes in mouse mobility (**Figure S1C**). Thus, our results indicate that any change in firing rate during SWDs could potentially be explained by the associated immobility and therefore cannot be attributed to SWDs *per se*. Collectively, these observations indicate that SWDs — surprisingly — are not defined by any meaningful changes in mean firing rates.

### Neuronal population synchrony and rhythmicity are profoundly elevated during SWDs

Unlike neuronal firing rates, the temporal organization of cortical firing is dramatically altered during SWDs. Pearson correlations of neuronal spike trains measure the degree of consistent co-activity in simultaneously recorded neurons and, therefore, can be used to resolve synchronous firing patterns within neuronal populations (see *Pearson correlations of neuronal spike trains* in STAR Methods). Indeed, we observed substantially increased synchrony among neurons, relative to baseline periods (**Figure 3A**). To quantify such synchrony, we first binned spike times (50ms bin) and then compared pairwise spike train correlations across non-ictal periods (“Baseline”) and SWD conditions (**Figure 3B**). From this analysis, we found elevated mean Pearson correlation coefficients (*R*) in S1 (*R* = +0.14, *n* = 13 recording sessions, *p* = 1.1×10^− 6^, paired t-test) and M2 (+0.06, *n* = 3, *p* = 0.03). Notably, Meyer et al. (2018) used a similar approach on indirect measures of neuronal spiking activity (i.e., 2-photon GCaMP-imaging) in V1 and surprisingly showed that mean Pearson correlations *decrease* during SWDs in the Stargazer model of absence epilepsy.

**Figure 3.**
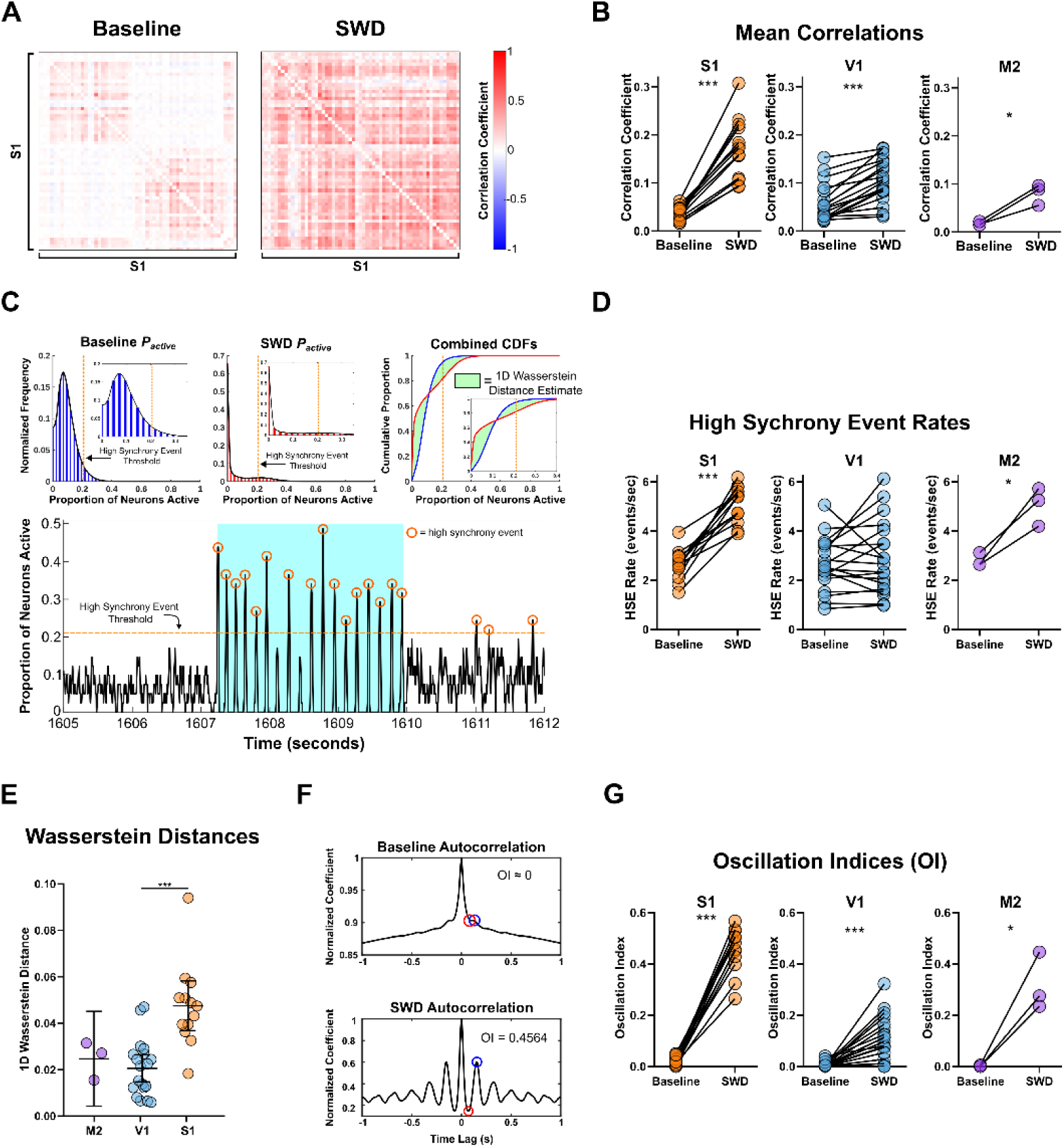
Synchrony and rhythmicity increase during SWDs. **(A)** Correlations during baseline and SWD from one recording session with neurons in S1. **(B)** Mean correlation coefficients during baseline and SWD. Each session is represented by a pair of connected circles. **(C)** Top: probability distributions of proportion of neurons active (***P****active*) within an entire example session. Also shown are the cumulative distribution functions (CDF) and the 1D Wasserstein distances between them (green shading). Insets show the same data with expanded x-axis. Bottom: an example of ***P****active* during 7 seconds of a session with the SWD highlighted with cyan shading. The orange dashed line in all plots corresponds to the 95th percentile score of the baseline ***P****active* distribution used as a threshold for detecting HSEs. Note the increased incidence of HSEs during the SWD. **(D)** HSE rates during baseline and SWD, following the same organization as in Panel B. **(E)** 1D Wasserstein distances between ***P****active* distributions. Each circle represents one recording session. **(F)** Example autocorrelograms of ***P****active* during baseline (top) and SWD (bottom). The oscillation index (OI) is the difference between the first peak at lag > 0 (blue circle) and minimum value between this first peak and lag = 0 (red circle). **(G)** OIs during baseline and SWD. * p < 0.05, ** p < 0.01, *** p < 10^−3^.

By contrast, using more direct measures of neuronal activity afforded by silicon probe recordings, we observed that correlated firing activity among V1 neurons is *elevated* during C3H/HeJ SWDs, relative to baseline periods (+0.05, *n* = 19, *p* = 7.6×10^−7^), albeit to a smaller extent than observed changes in S1 (**Figure 3B**). Because these correlations are sensitive to bin size, we repeated these comparisons usings parametrically varied bin durations from 10ms to 210ms (**Figure S2A**) and found that mean correlation changes are consistent up to bin durations of ∼150ms, a value approaching the period of a single SWD cycle (∼167ms). With such long bin durations, multiple SWD cycles are often combined into single bins, thereby obscuring the fine-scale temporal organization of population-level activity.

In addition to Pearson correlations, synchrony can also be quantified as the proportion of simultaneously active neurons within a defined time window. Thus, we discretized neuronal spike trains into 25ms time bins and identified the proportion of neurons with at least one spike per bin (***P****_active_*). During baseline periods, ***P****_active_* remains consistently low over time. During SWDs, however, ***P****_active_* alternates between very low and very high values, reflecting the rhythmic alternations between neuronal silence and population bursts that occur during SWDs (**Figure 3C**). We defined peaks in ***P****_active_* as high synchrony events (“HSEs”, see *Detection of HSEs* in STAR Methods) and computed the rate at which these events occur during baseline and SWD periods. HSE rates increased during SWDs in S1 (+2.5 HSE/s, *n* = 13 recording sessions, *p* = 5.5×10^−7^, paired t-test) and M2 (+2.3 HSE/s, *n* = 3, *p* = 0.04, **Figure 3D**). The HSE rate in V1 did not change during SWDs (+0.3 HSE/s, *n* = 19, *p* = 0.33, **Figure 3D**). These results remained robust across a range of bin durations (**Figure S2B**), but effect sizes were generally found to be largest using a 30ms bin. Thus, we conclude that SWDs are generally associated with an increase, not a decrease, in correlated neural activity patterns. In addition to measuring changes in HSE rates between baseline and SWD periods, we also used HSEs to estimate the number of neurons active during single SWD cycles. Using the mean HSE magnitude (i.e., the mean proportion of neurons active during HSEs) from each recording session as a sample, we found that HSEs recruit between 20-40% of local neurons during SWDs (**Figure S2C**). This finding represents the first explicit estimate of neuronal population recruitment during single SWD cycles.

HSEs represent brief windows of elevated population activity. However, SWDs are also characterized by periods of near-complete neuronal silence. To evaluate overall changes in the temporal organization of population activity — both elevated population spiking and silence — we measured the overall distance between distributions of ***P****_active_* by computing the 1D Wasserstein distance between baseline and SWD distributions (see green shading in **Figure 3C**). This distance measures overall differences in ***P****_active_* distributions and is sensitive to both synchronous silence and spiking. Therefore, we consider the measure a relatively unbiased assessment of the extent of change in neuronal population activity during SWDs. One-way ANOVA measures revealed a significant effect of brain region on 1D Wasserstein distances (F = 9.3, *p* = 4.1×10^−6^), and such distances in S1 were greater than those in V1 (*p* < 0.01, Bonferroni-corrected *post hoc* tests). Thus, SWDs more profoundly alter population-level neuronal activity in S1 than in V1.

Finally, in addition to becoming more synchronous, the population-level activity of neurons also becomes highly rhythmic during SWDs. To quantify changes in rhythmicity, we first calculated the autocorrelation of ***P****_active_* during baseline and SWD periods (see *Quantifying rhythmicity of population activity* in STAR Methods, **Figure 3F**). We then computed an oscillation index (OI, Kleiman-Weiner et al., 2009) from the resultant autocorrelograms. OI values increased from baseline to SWD in S1 (+0.44, *n* = 13, *p* = 4.0×10^−10^, paired t-test), M2 (+0.32, *n* = 3, *p* = 0.037), and V1 (+0.10, *n* = 19, *p* = 9.0×10-5) (**Figure 3G**). We also observed significant SWD-associated differences in correlated activity, HSEs, OI values and Wasserstein distances among caudoputamen and hippocampal neuronal populations (**Figure S3**).

### Cortical neurons are highly phase-locked to SWDs

In total, 2,839 SWDs and 2,228 isolated neurons (**Table S2**) were recorded. The spike times of many cortical neurons were phase-locked to or near SWD troughs (**Figure 4A**). To quantify phase-locking, SWDs were first segmented into individual cycles wherein the most negative amplitude of two successive troughs defined the start and end of a cycle. Neuronal spikes were then assigned a phase value between 0° and 360° (**Figure 4B**). Mean vectors for each neuron were computed by taking the complex sum of action potential phases (see *Spike-SWD phase-locking* in STAR methods for details, **Figure 4C**). Magnitude of phase-locking was estimated by the mean vector length of all neurons in that region (**Figure 4D,E**). S1 had the longest mean vector length (0.66), followed by M2 (0.54), and V1 (0.23) (**Figure 4E**). A linear mixed-effects model with session as a random effect revealed that mean vector lengths in all regions significantly differed from one another (*p* < 0.01, Tukey-adjusted pairwise comparisons of estimated marginal means, see **Table 1** for all comparisons), indicating that the magnitude of phase-locking varies among different brain regions. We also determined the percentage of neurons in each brain region that exhibited significant phase-locking. To this end, for each neuron, spike phases were randomly shifted 100,000 times, and vector lengths were calculated to generate a null distribution (**Figure S4**). Only neurons with observed vector lengths greater than nearly all the null distributions were considered significantly phase-locked (see STAR methods for details). S1 had the highest percentage of significantly phase-locked neurons (86%, 675/785), followed by M2 (85%, 298/350), and V1 (54%, 275/513) (**Figure 4E**). By contrast to vector lengths, vector angles yield insight into the timing of neuronal spiking within SWD cycles. Considering only significantly phase-locked neurons, S1 showed the earliest mean vector angles (-17.9°), followed in order by M2 (-12.9°), and V1 (31.1°) (**Figure 4F**). S1 angles were significantly earlier than all regions other than M2 (*p* < 0.05, Tukey-adjusted pairwise comparisons of estimated marginal means, see **Table 1** for all comparisons).

**Figure 4.**
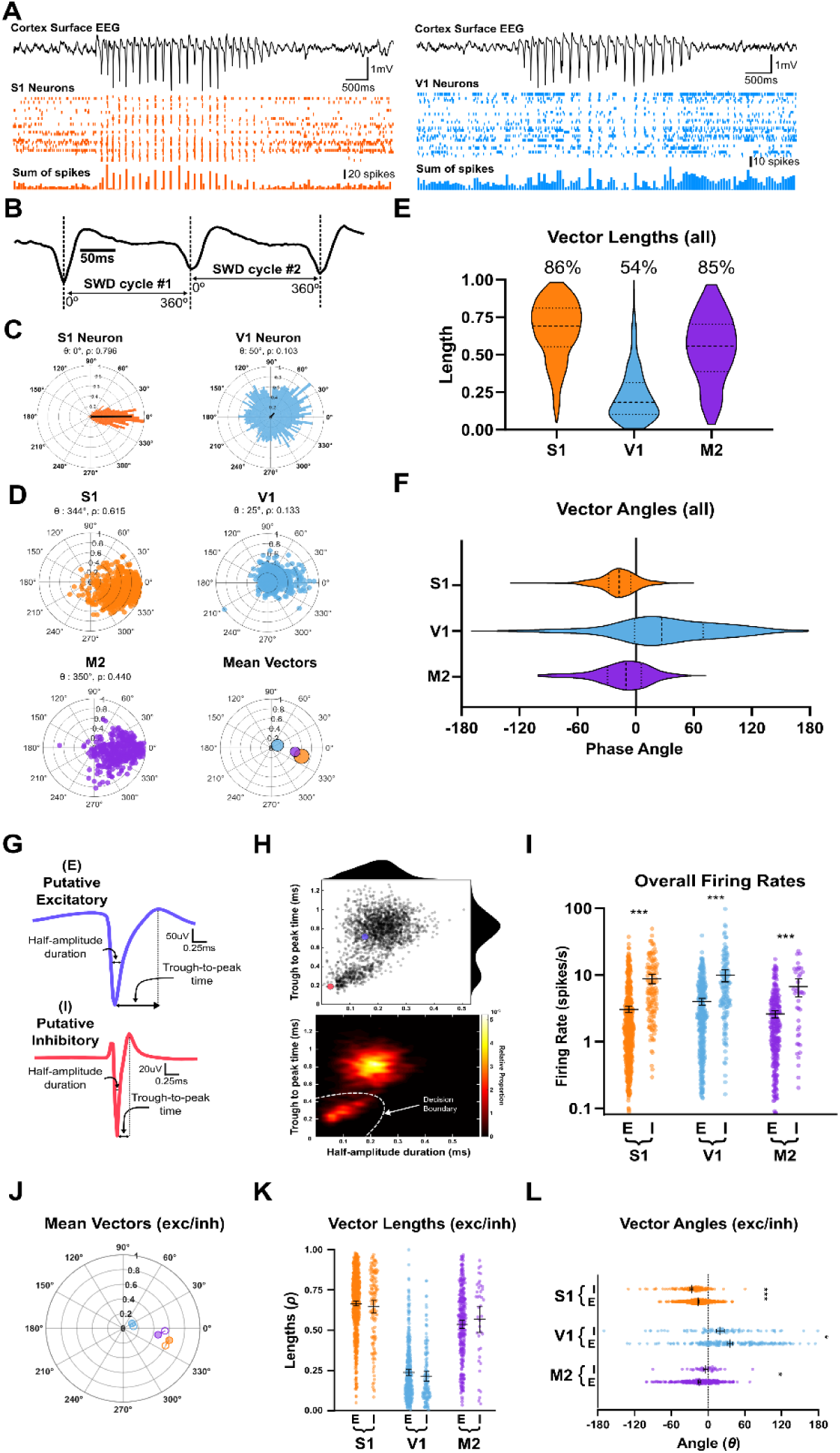
Neuronal activity is phase-locked to SWD troughs. **(A)** Left: Simultaneous recording of surface EEG and spike rasters for 32 neurons recorded in S1 during one SWD. Bottom bars show the summed spike count from all neurons. Right: Spike rasters for 37 V1 neurons during one SWD. **(B)** Surface EEG trace demonstrating how SWDs were divided into cycles. Vertical dashed lines indicate the cycle start/end. **(C)** Polar histograms of two example neurons, one in S1 showing strong phase-locking and another in V1 showing weak phase-locking to SWDs. Black lines represent the mean vector for that neuron. *θ* is the mean vector angle. *ρ* is the mean vector length. **(D)** Mean vectors of all neurons recorded in each brain region plotted separately. *θ* and *ρ* values here represent the mean vector angles and lengths across all neurons in that region and are plotted for each structure together in the bottom right polar plot. Circle size corresponds to the number of neurons recorded in that region. **(E)** Distributions of vector lengths in each brain structure. Middle dashed lines represent the median. Upper and lower dotted lines represent top and bottom quartiles. Vector lengths of all brain structures significantly differed from one another (see **Table 1**). Values above distributions indicate the percentages of neurons in each region that were significantly phase-locked (see **Figure S4**). **(F)** Distributions of vector angles for significantly phase-locked neurons in each region (**Table 1** shows all pairwise comparisons). **(G)** Two example spike waveforms from different neurons, one putative excitatory, one inhibitory, showing the two features used for classification: half-amplitude duration and trough-to-peak time. **(H)** Top: Half-amplitude durations and trough-to-peak times plotted for all neurons combined from all recordings. One-dimensional distributions are shown outside the top and right edges of the scatter plot. Enlarged scatter points in color correspond to the two example waveforms shown in panel **A**. Bottom: 2D histogram showing decision boundary derived from a mixture of Gaussians overlaid on the distribution of waveform features. **(I)** Overall firing rates of putative excitatory and inhibitory neurons. **(J)** Mean spike-SWD vector separated by putative neuron type. Open circles represent mean vectors for putative inhibitory neurons whereas shaded circles are those of excitatory neurons. Putative inhibitory neurons have earlier mean vector angles in all regions, except M2. **(K)** Vector lengths of putative excitatory and inhibitory neurons. **(L)** Vector angles of putative excitatory and inhibitory neurons. Bars indicate the 95% confidence interval.

**Table 1.**
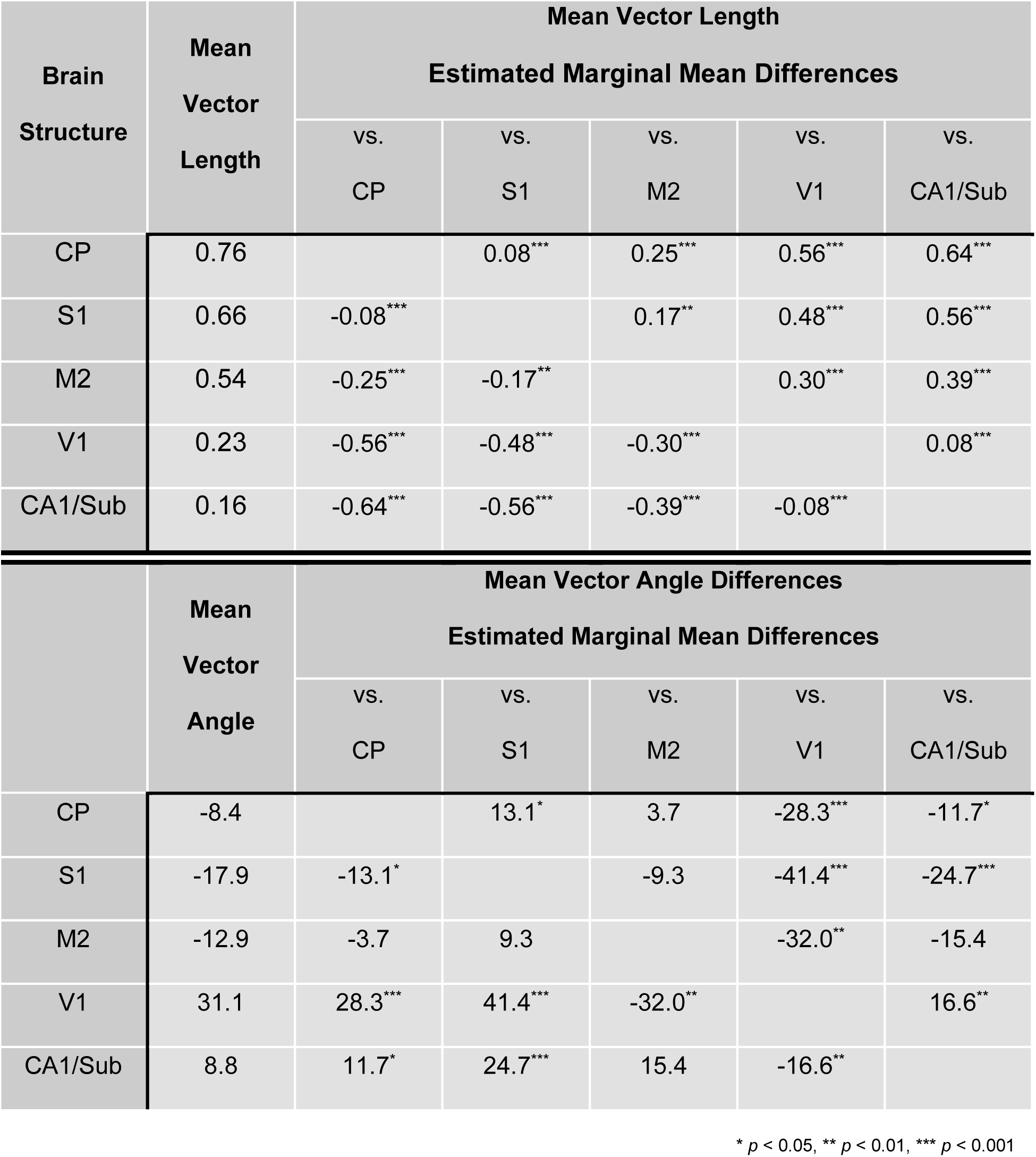
Mean vector lengths and angles by brain structure.

The observation that many neurons in all recorded cortical regions showed significant phase-locking suggests that SWDs engage widespread cortical networks. Moreover, mean vector angles in all regions are highly concentrated within a ∼60° range (-20 to +40°) surrounding SWD troughs. This ∼30ms active window represents just 1/6 of the full ∼167ms cycle period — a remarkably brief burst of coordinated activity across distributed structures. Finally, our finding that S1 neurons show strong phase-locking and the earliest vector angles accords with its proposed role as a trigger zone for SWDs (Meeren et al., 2002).

Next, we aimed to resolve the extent of neuronal engagement during SWDs. To this end, we calculated a *participation proportion* for each significantly phase-locked neuron. We classified a neuron as participating if it spiked at least once during a specific interval, defined either as an entire SWD (“per SWD”) or a 60ms window centered on SWD troughs (“per cycle”). To limit the contamination of unrelated firing from non-phase-locked neurons, only significantly phase-locked neurons were included in this analysis. *Per SWD* participation proportions averaged between 0.83 and 0.92 across brain regions (i.e., neurons spiked at least once during 83-92% of all SWDs). These participation proportions followed highly skewed distributions with most neurons active during most SWDs (*p* <0.001 for D’Agostino K^2^ tests in all regions, **Figure S5A**, **Table S3**). However, *per cycle* participation proportions also had skewed distributions (*p* < 0.05) wherein means ranged from 26 to 33% (**Figure S5B**, Table S3). This conclusion is consistent with the 20-40% estimate derived from our HSE analyses (see **Figure S2B** and **Table S2**). Thus, most neurons spike at least once during most SWDs, but very few neurons spike on every SWD cycle. These results reflect two basic statistical features of neuronal activity during SWDs and, surprisingly, indicate that SWDs likely reflect an emergent property that involves a substantial population of only modestly engaged neurons. That is, SWDs do not appear to involve significant populations of highly active and highly engaged neurons.

### Putative inhibitory neurons fire early during a SWD cycle

In addition to varying by brain region, we also determined whether phase-locking measures vary as a function of neuronal type. We were specifically interested in understanding the basis for the sustained neuronal silence that occupies nearly 5/6 of a SWD cycle (**Figure 4C,D**) and speculated that prolonged inhibitory neuron activity may be responsible. To characterize the activity of inhibitory neurons, we leveraged differences in action potential waveform shapes to distinguish inhibitory neurons from other neuron types (**Figure 4G**). Using half-amplitude duration and trough-to-peak time as features (Barthó et al., 2004), two distinct waveform clusters are apparent and can be separated using a decision boundary derived from a mixture of Gaussians (see *Classification of spike waveforms* in STAR Methods, **Figure 4H**). As expected, baseline firing rates in putative inhibitory neurons were higher than excitatory neurons in all brain regions (*p* < 0.001, unpaired t-tests, **Figure 4I**), reflecting the standard properties of the predominant fast-spiking GABAergic interneuron subtype. We also observed that putative inhibitory neurons were strongly phase-locked to the SWD (**Figure 4J**) and their vector lengths did not differ from putative excitatory neurons (*p* > 0.05, **Figure 4K**). Surprisingly, the vector angles of putative inhibitory neurons were significantly earlier in S1 and V1 (*p* < 0.05 for all regions, **Figure 4L**). In M2, however, putative inhibitory neurons lagged putative excitatory neurons (i.e., phase angles were delayed, *p* < 0.05). Similarly, baseline firing rates in putative inhibitory neurons were higher than those of putative excitatory neurons in both CP and CA1/Sub (p < 0.001, **Figure S6A**). In these subcortical structures, putative inhibitory and excitatory neurons exhibited similar phase-locking strengths (**Figure S6B,D**). However, putative inhibitory neurons in CP fired significantly earlier during the SWD cycle than putative excitatory neurons, whereas no difference in vector lengths was observed in CA1/Sub (**Figure S6C**). In summary, putative inhibitory and excitatory neurons are equally phase-locked to SWD cycles, and inhibitory spiking occurs earlier during an SWD cycle than excitatory neuron spiking in both S1 and V1. We found no evidence of sustained inhibitory neuron spiking during the long silent phase of SWD cycles. Thus, it remains plausible that a brief burst of intermixed inhibitory and excitatory neuron activity may lead to overall population silence. If true, then such a reduction would likely *not* be driven by ionotropic, GABA_A_ receptor-mediated inhibition.

### The order of neuronal population activity is conserved across SWDs

Thus far, we observed that SWDs appear to be supported by many weakly engaged neurons, and that putative inhibitory neurons generally fire earlier than excitatory neurons during a SWD cycle. Next, we aimed to resolve whether individual cortical neurons — despite only weak participation during SWDs — follow defined temporal sequences during seizures, as they do during sleep and anesthesia (Luczak et al., 2007), or whether the activity of such neurons is essentially stochastic. As SWD-associated firing among individual cortical neurons was sparse, it was difficult to resolve consistent, temporally ordered ensembles among recorded neurons during individual SWDs, as one can during single memory-associated sharp-wave ripples (Diba & Buzsáki, 2007; Pfeiffer & Foster, 2013). Nonetheless, we were able to quantify the temporal consistency with which individual neurons fired within and across successive SWDs. We were specifically interested in understanding whether a single neuron’s temporal position within an SWD cycle is fixed across multiple SWDs and, if a neuron fires multiple times during an SWD, whether it’s temporal position within a SWD cycle is also fixed. To this end, we examined the relative firing times of significantly phase-locked cortical neurons simultaneously recorded in a single recording session by creating peri-event time histograms (PETHs) centered on SWD troughs and displayed as a gray-scale heat map (**Figure 5A**). Visualizing activity in this manner revealed that individual cortical neurons fire in consistent temporal positions within a SWD cycle, and that sequential patterns clearly emerge when evaluating SWDs in aggregate within a single recording session. (Importantly, because we could not reliably identify consistent neuronal groups across SWDs, we do not classify these activity patterns as ensembles in the strict sense.) To further evaluate such temporal consistency, we first separated SWDs into two groups (i.e. first half and second half) and then separately computed the center-of-mass of trough-centered PETHs for each group (see *Sequential activity in SWD trough-centered PETHs* in STAR Methods). The red line overlayed on **Figure 5A** indicates the center-of-mass of cortical neurons during first half SWDs, whereas the green open circles indicate their corresponding center-of-mass during the second half SWDs. **Figure 5B** displays the two centers-of-mass for three individual neurons indicated in red, blue, and yellow circles in **Figure 5A**. We then compared the sequence of centers-of-mass in the first half to the sequences from the second half using Spearman’s rank correlations to determine if sequences were conserved (**Figure 5C**). For this analysis, rank correlations derived from neurons on a single shank of the silicon probe were considered unique observations. After excluding shanks and/or recordings with too few significantly phase-locked neurons, the percentage of eligible observations showing significant preservation of temporal order was 76% in S1 (19/25), 83% in M2 (5/6), and 35% (7/20) in V1 (*p* < 0.05, Spearman’s rank correlations with Bonferroni adjustment applied per brain region, **Table 2**).

**Figure 5.**
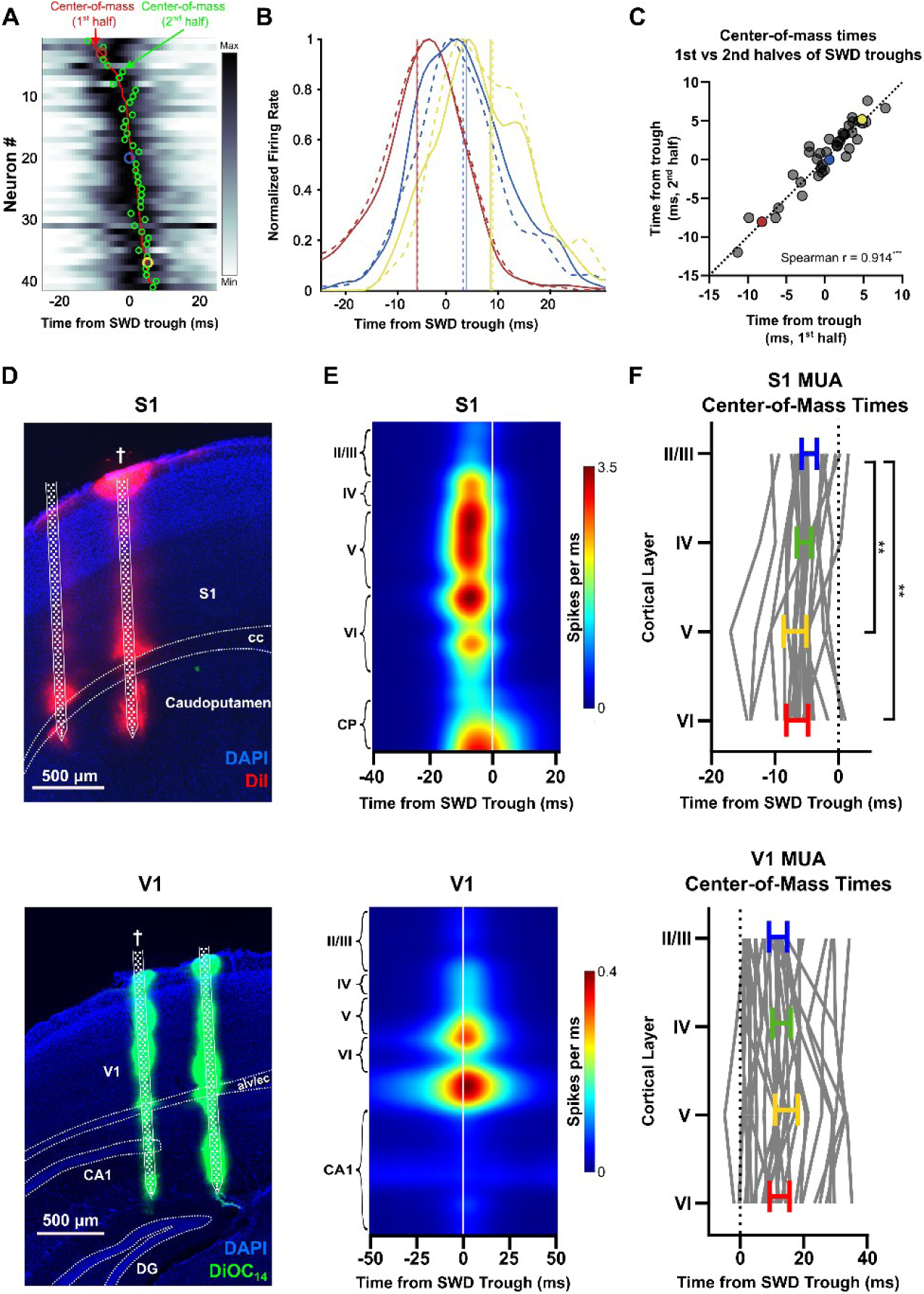
Sequential activation of cortical neurons around SWD troughs. **(A)** SWD trough-centered PETH sorted by center-of-mass times. Each SWD was divided into a first and second half. The red line indicates the center-of-mass times derived from the 1^st^ halves of all SWDs in the recording, whereas the open green circles indicate center-of-mass times derived from the 2^nd^ halves of all SWDs (e.g., cycles 1-7 versus 8-14 in a 14-cycle SWD). Note the sequential order of the centers-of-mass and the conservation of that order from the 1^st^ to 2^nd^ half of SWDs. Open circles in red, blue, and yellow are data from the same neurons show in panels B and C. **(B)** SWD trough-centered PETHs from three different neurons. Solid lines are PETHs derived from the 1^st^ halves of all SWDs and dotted lines are PETHs derived from their respective 2^nd^ halves. Vertical lines indicate the center-of-mass for PETHs of corresponding colors and line styles. **(C)** Scatter plot of center-of-mass times derived from all 1^st^ versus all 2^nd^ halves of SWDs. Dotted diagonal line is the identity line. *** p < 10^−3^. **(D)** Representative histology showing fluorescent dye from probe tracks in S1 (top) and V1 (bottom). † indicates shank used for MUA plots in Panel B. **(E)** Average MUA PETHs from vertically sorted electrodes and cortical layers labeled. Note the difference in firing rate (color scale) limits and time (x-axis) limits between S1 (top) and V1 (bottom). **(F)** Center-of-mass times relative to SWD troughs. Colored horizontal lines and bars indicate the means and 95% confidence intervals. Gray lines indicate center-of-mass times from single shanks from all included recordings. Note the difference in time (x-axis) scale between S1 (top) and V1 (bottom). ** p < 0.01

**Table 2.** Spearman rank-order correlations per brain structure.

| Brain Region | # observations with significant Spearman's r | # eligible observations | Percentage significant |
| --- | --- | --- | --- |
| CP | 6 | 13 | 46.2% |
| M2 | 5 | 6 | 83.3% |
| CA1/Sub | 2 | 7 | 28.6% |
| S1 | 19 | 25 | 76.0% |
| V1 | 7 | 20 | 35.0% |

The simplest explanation for the observed temporal consistency is that these neurons reside in different cortical layers, and that their firing sequence reflects the sequential engagement of cortex by the SWD (Polack et al., 2007). We therefore compared the multi-unit activity (MUA) around SWD troughs across cortical layers. We used MUA to estimate the overall activity within a cortical layer rather than the mean activity of isolable single units because MUA provides a more robust measure of the overall activity of a local population of neurons. We computed the peri-trough PETHs on each electrode, averaged them from electrodes within the same cortical layer, and computed their center-of-mass times (see *Multi-unit activity (MUA) for laminar activation analysis* in STAR Methods, **Figure 5D**). A stereotyped temporal pattern of activation across cortical layers was clearly evident in S1. One-way repeated measures ANOVA revealed a significant effect of cortical layer on timing (F = 10.4, *p* = 0.002) with layers V and VI significantly preceding layer II/III (*p* < 0.05, Bonferonni-corrected *post hoc* tests (**Figure 5F**). By contrast, no such temporal order was evident in V1 (F = 2.1, *p* = 0.1) and we found no differences between individual layers (*p* > 0.05 for all pairwise comparisons). Considering that neuronal activity within layers V and VI tend to lead other layers during an SWD cycle, it is likely that the temporally ordered sequences we observed do, at least in part, reflect cortical layers activating at different times.

### 6Hz and 100Hz electrical stimulation of S1 and V1 can both induce and terminate SWDs

The observed phase-locking and synchrony among cortical neurons indicate that SWDs engage both S1 and V1, but that S1 recruitment by SWDs is substantially stronger. This conclusion is consistent with the longstanding view that S1 plays an outsized role in SWD generation. We therefore hypothesized that SWDs are more sensitive to manipulations of S1 than V1. To test this hypothesis, we implanted twisted electrode pairs bilaterally in either S1 or V1 and stimulated each region using two distinct patterns: (1) 6Hz burst stimulation designed to induce SWDs (**Figure 6A**), and (2) 100Hz continuous stimulation designed to interrupt ongoing SWDs (**Figure 6C**). A linear mixed effects model with stimulation current intensity as a fixed effect, and mouse identity/session as a random effect, revealed that 6Hz stimulation of either S1 or V1 induced SWDs in both regions at all current intensities except 10 µA (*p* < 0.01, **Figure 6B**, Dunnett’s *post hoc* tests relative to 0µA baseline). Using a linear mixed effects model with the same design, we also found that 100Hz stimulation of either S1 or V1 terminated ongoing SWDs at current intensities greater than except 10 µA (*p* < 0.01, **Figure 6D**). Considering that S1 and V1 were capable of both inducing and terminating SWDs, and that phase-locking and synchrony are elevated in many brain regions (**Figures 1** and **6**), we conclude that SWDs involve widespread brain networks capable of initiating and maintaining SWDs.

**Figure 6.**
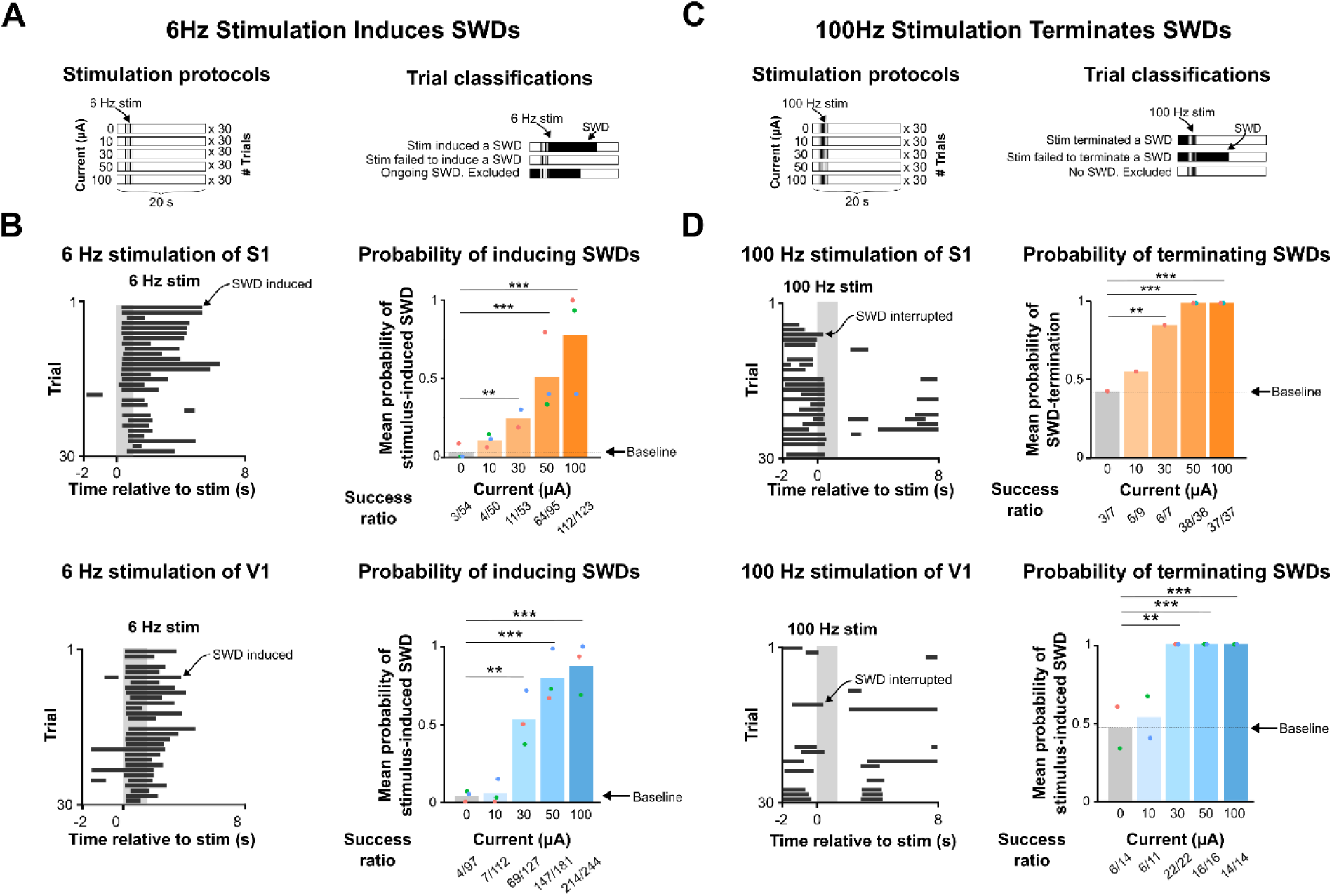
S1 and V1 stimulation induces or terminates SWDs at 6Hz or 100Hz, respectively. **(A)** Stimulation protocol for inducing SWDs with 6Hz stimulus. For SWD induction, trials were classified as successful if stimulation induced an SWD or unsuccessful if not. Trials with already ongoing SWDs during stimulation were excluded. **(B)** Left: example 6Hz stimulation session in S1 (top) and V1 (bottom). Gray shading indicates 6Hz stimulation duration. Horizontal black bars represent SWDs. Right: probability of SWD induction plotted against different levels of current intensity in S1 (top, orange) and V1 (bottom, blue). **(C)** Stimulation protocol for terminating SWDs with 100Hz stimulus. Trials were classified as successful only when ongoing SWDs were terminated during the 100Hz stimulus duration. Trials without ongoing SWDs during stimulation were excluded. **(D)** Same as Panel B but for 100Hz stimulus terminating ongoing SWDs. ** p < 0.01, *** p < 10^−3^.

## DISCUSSION

Here, we set out to resolve the ubiquity and magnitude of cortical engagement during SWDs, the electrographic hallmark of absence seizures. The data presented here support the conclusion that SWDs engage widespread areas of cortex (Gibbs et al., 1935), but that significant regional differences exist. We demonstrate that S1 cortex shows stronger phase-locking, more synchrony, and enhanced oscillatory activity during SWDs, relative to M2 and V1. S1 was also engaged earlier in the SWD cycle than all other recorded structures. Thus, our observation that S1 is highly engaged and active early during an ongoing SWD aligns with previous work implicating S1 as an important, SWD-generating region (Atherton et al., 2023; Meeren et al., 2002; Polack et al., 2007).

Whereas neuronal activity in S1 showed an expected phase-locking, synchrony, and rhythmicity during SWDs, our V1 recordings contradict previous findings. By employing 2-photon imaging of GCaMP-expressing cortical neurons in the Stargazer model of absence epilepsy, Meyer et al. (2018) reported that V1 neuronal activity during SWDs is characterized by reduced synchrony. By contrast, we found that V1 neurons in the C3H/HeJ mouse are clearly synchronously active during SWDs, albeit less so than in other cortical regions (i.e., S1 and M2). First, we found a small, but significant, increase in spike train correlations in V1. Specifically, in each of the 19 V1 recording sessions included in our dataset, spike train correlations increased during SWDs (**Figure 3B**). We also report that a substantial fraction of V1 neurons (54%) are significantly phase-locked to ongoing SWD activity (**Figure 4E**), and that population-level activity becomes increasingly rhythmic during SWDs (**Figure 3G**), indicating some level of neuronal entrainment. Moreover, V1 spike rasters (e.g., **Figure 4A**) reveal clear population-level synchrony during SWDs, at least during a subset of cycles. Regarding the extent of V1 engagement during SWDs, we attribute the discrepancy between our findings and those of Meyer et al. (2018) primarily to differences in recording methodology. Using electrophysiological recording techniques sampled at 30kHz, our results are derived from sub-millisecond sampling of neuronal spike times and, therefore, can capture the temporal structure of neuronal activity with high resolution. By contrast, the two-photon Ca^2+^-imaging techniques utilized by Meyer et al. (2018) relied on a sampling frequency of 20Hz, leading to a the relatively prolonged sampling interval of 50ms, a value roughly 1/3 the duration of a single SWD cycle. Further restricting the temporal resolution of GCaMP-based imaging approaches are the slow rise (*τ* ≈ 50ms) and decay (*τ* ≈ 300ms) kinetics of the GCaMP6m sensor. Given limitations on the temporal resolution of two-photon GCaMP-imaging, we argue that the extracellular electrophysiology approach provides a more accurate estimate of pairwise neuronal spike train correlations.

Nonetheless, in accordance with Meyer et al. (2018) and McCafferty et al. (2023), we also found that firing rates generally decreased during SWDs and expanded the finding to include M2 (**Figures 2** and **S1**). However, because SWDs are accompanied by behavioral arrest (Blumenfeld, 2005a), we asked whether the firing rate changes we observed during SWDs simply reflect immobility rather than the seizure itself. Across all recorded cortical structures (S1, V1, M2), firing rates were reduced during both SWDs and non-SWD immobile states, relative to mobile states, and linear regression showed that immobility accounted for the majority of SWD-related firing rate changes (R² = 0.57–0.88). This relationship extended to subcortical structures as well, including CP and CA1/subiculum (**Figure S1**). Together, these findings indicate that mean firing rate changes during SWDs are largely attributable to concurrent immobility rather than to the seizure state per se, suggesting that SWDs are not fundamentally defined by altered mean firing rates. Notably, Buzsaki et al. (1988) speculated that immobility was a necessary condition for SWD generation. Support for this hypothesis has been recently provided insofar SWDs were abolished when subjects were in high-arousal states asked to actively attend (McCafferty et al., 2023) or forced to ambulate (Dong et al., 2024). If overt neuronal hyperexcitability does not initiate or maintain SWDs, and behavioral arrest is required for SWD formation, then a central question becomes: how do immobility and/or decreased arousal promote regional neuronal hypersynchrony that appears to initiate and sustain SWDs? In the absence of overt hyperexcitability, we speculate that absence seizures primarily reflect changes in the organization of neuronal population activity, and not the magnitude of activity. If true, then pharmacological therapies for absence epilepsy may be misguided when they specifically target neuronal hyperexcitability. Indeed, roughly one third of absence epilepsy patients are pharmacoresistant (Cnaan et al., 2017).

SWD-induced changes in the organization of population activity are clearly apparent when examining the proportion of neurons firing at least once within a moving time window (“***P****active*” in **Figure 3C**). ***P****active* oscillates between high and low values during the “spike” and “wave” phase of SWDs, reflecting population activity and silence, respectively (**Figure 3C**). We refer to high magnitude peaks in ***P****active* as “high synchrony events” (HSEs in **Figure 3D**), and HSEs generally represent the largest events we observed. HSEs vary slightly in magnitude by brain structure, but on average, ∼30% of recorded neurons spike during individual cycles of SWDs (**Figure 2C**), a value that is also consistent with our finding that, on average, any given neuron participates in ∼30% of SWDs (**Figure S5B**). While estimates have been made for other large population events, such as sharp wave-ripples in the hippocampus (Buzsáki, 2015) and neocortical UP states (Cossart et al., 2003), our analysis marks the first explicit attempt to estimate the proportion of neurons participating during a SWD.

Similar to the population-level activity that occurs during the SWD “spike”, the silence during the “wave” component of the SWD is also profound. The prolonged neuronal silence often observed in the thalamus during SWDs is thought to be achieved by powerful inhibitory drive from the reticular thalamus (Steriade, 2005). However, as neurons of the reticular thalamus do not project to the cortex, the mechanism(s) responsible for prolonged cortical silence remain unknown. Here, we present near-complete suppression of activity during the wave component in S1 and M2, even among putative inhibitory interneurons (**Figure 4**). This observation suggests that the prolonged cortical silence is not sustained by local interneurons that remain active during the wave, unless such local inhibition recruits the activation of metabotropic GABAB receptors that have a kinetic profile that closely aligns with the period of SWD cycles (Beenhakker & Huguenard, 2010; Bettler et al., 2004; Destexhe, 1998; Lu et al., 2020). If these receptors are not recruited during the wave portion of the SWD, the prolonged silence likely reflects either a loss of excitatory drive from thalamic structures, an intrinsic property of cortical neurons that limits sustained activation, or some combination of the two.

In summary, we found that SWDs reflect population-level hypersynchrony and that such hypersynchrony exists in several cortical areas, including V1, which was previously reported to desynchronize during SWDs. We also argue that the organization of population activity, specifically the synchronized firing of many neurons, is what truly typifies SWDs, not simple changes in firing rates. Finally, we show that SWDs begin only after immobility has already set in and propose that immobility-driven firing-rate reductions establish an SWD-permissive state rather than reflecting SWD activity itself. Collectively, these findings advance a more complete understanding of SWDs and establish a new perspective for the development of future treatments for absence epilepsy.

## Materials availability

Custom-made 3D-printed recording chamber design files (**Figure S7**) can be found here: https://github.com/scottkil/SiProbe_RecordingChamber

## Data and code availability

Data and relevant code for analysis can be found here: https://github.com/scottkil/PSR.

## STAR★METHODS

### EXPERIMENTAL MODEL AND STUDY PARTICIPANT DETAILS

#### Mice

Female C3H/HeJ mice (Strain #:000659) aged 6-10 weeks were purchased from The Jackson Laboratory and used for the experiments described here. Mice were housed at 23-25°C under an artificial 12-hour light-dark cycle with food and water *ad libitum*. All procedures were approved by the University of Virginia Animal Care and Use Committee (Charlottesville, VA, USA) and performed in accordance with the NIH Guide for the Care and Use of Laboratory Animals.

### METHOD DETAILS

#### Head-fixation and surface EEG electrode implant

Mice were anesthetized with vaporized isoflurane (2-4% for induction, 1-2% for maintenance) mixed with pure oxygen and delivered at a rate of 1 liter/min. Body temperature was maintained at 37°C with a temperature probe and thermostatic heating pad placed underneath the mouse. Lubricating ointment was applied to the eyes. The scalp was first shaved, and then depilatory cream was applied to further remove hair. The scalp was then cleaned with alternating scrubs of betadine and 70% ethanol three times. After making a sagittal incision along the midline, the underlying periosteum was scrubbed away with sterile cotton tipped applicators, and an applicator dipped in 3% hydrogen peroxide was used to clean the skull and increase the visibility of cranial sutures. Once the dorsal surface of the skull was fully cleared of periosteum, custom-designed titanium headbars (Star Rapid, China) were fixed to the posterolateral edges of the parietal bones with dental cement (C&B Metabond, Parkell, New York, USA).

Next, a burr hole was made in the frontal bone over the secondary motor cortex for a surface EEG electrode. Two additional burr holes were made at the posterior aspect of the interparietal bone overlying the cerebellum, lateral to the midline, for reference and ground electrodes. A custom electrode array, consisting of insulated stainless-steel wires (A-M Systems, Washington, USA) soldered to a 3-position header of connector pins (Digikey, Minnesota, USA), was lowered into the surgical field. The wires were carefully placed under the skull in the burr holes using forceps and fixed in place with UV-curing glue (Wave, SDI, Illinois, USA). After all wires were glued, the electrode array was cemented to the posterior edge of the skull with care taken to avoid covering the remaining exposed skull with dental cement. Opaque silicone elastomer (World Precision Instruments, Florida, USA) was then applied to the remaining exposed skull and allowed to cure for five minutes. Ketoprofen (5mg/kg) was administered peri-operatively to reduce pain and inflammation. Mice recovered under daily monitoring in their home cage for seven days before entering the next stage of the experiment.

#### Acute silicon probe recordings

Mice were habituated to the head-fixation apparatus for a minimum of three one-hour sessions. During these sessions, mice were head-fixed to a stationary rod on top of a polystyrene ball, which could be rotated forwards and backwards by ambulation. After the habituation phase, one day prior to the acute silicon probe recording, a craniotomy was performed over the recording site, and a protective recording chamber was installed. Briefly, the silicone elastomer on the skull was peeled away and a rectangular (∼1.5 x ∼1.5mm) craniotomy was made in a mouse under isoflurane anesthesia. The following coordinates with respect to bregma were used as center points for craniotomies: S1 - AP: 0mm, ML: -3mm; V1 - AP: -3mm, ML: -3mm; M2 - AP: +2.3mm, ML: -1.5mm. The craniotomy was made by repeatedly tracing a rectangular outline with a dental drill, removing a thin layer of skull with each pass, and rinsing the site with cooled artificial cerebrospinal fluid (ACSF) between passes. When the fragment within the outline had been sufficiently loosened from the surrounding skull, it was soaked in ACSF for five minutes to soften any remaining thinned bone and then carefully lifted away from the brain with forceps. A custom-designed 3D-printed protective recording chamber (**Figure S7**) was then centered on the craniotomy and cemented to the skull. An Ag/AgCl ground wire was fixed to the skull within the chamber with UV-curing glue. Finally, the chamber was filled halfway with triple antibiotic ointment and sealed with silicone elastomer to retain moisture overnight.

Just prior to the acute silicon probe recording sessions, fluorescent dyes — 2mg/mL DiI (Biotium, California, USA) or DiOC14 (Biotium, California, USA) dissolved in 100% EtOH — were applied to the probes. Probes were positioned horizontally in a micromanipulator with electrodes facing down. A vertically oriented syringe with a droplet of dye solution hanging from the tip was lowered onto the probes. Probes remained in the droplet for 5 minutes. During the recording sessions, mice were head-fixed to the recording apparatus as they were during habituation sessions. Using a long working distance digital microscope (Dino-Lite, California, USA) to visualize the area within the recording chamber, the silicone elastomer was peeled away with forceps, and any remaining ointment was rinsed out with ACSF. Once the brain was clearly visible, the silicon probes were manually moved to the brain surface. Then, the probes were lowered into the brain at a constant rate of 2µm/s with a motorized micromanipulator system (Sutter Instrument, California, USA). The recording chamber was filled halfway with ACSF and then topped with mineral oil to minimize evaporation and dehydration of the brain during the recording.

Upon reaching the target depth, probes were allowed to settle for an additional 30 minutes before recording commenced. Data from the silicone probe electrodes was amplified using two 128-channel RHD headstages (Intan technologies, New York, USA) and digitized at 30kHz.

Analog input channels for the surface EEG and TTLs from the rotary encoder were also sampled at 30kHz. Surface EEG was amplified 5,000 times and filtered between 1 - 1kHz with a differential amplifier (A-M Systems, Washington, USA).

After the recording session, the probe was gently raised out of the brain at a rate of 10µm/s and set aside. The recording chamber was cleaned with sterile cotton-tipped applicators and ACSF rinses. The chamber was then filled with triple antibiotic ointment and sealed with silicone elastomer. The mouse was then removed from the head-fixation apparatus and returned to the home cage. Silicon probes were rinsed with deionized water and ethanol before being soaked in 0.05% Trypsin-EDTA (Thermo Fisher Scientific, Massachusetts, USA) for at least 30 minutes, followed by 1% Tergazyme (Alconox, New York, USA) for an additional 30 minutes. Both soaks were performed in solutions at 60°C and followed by a final water rinse.

#### Euthanasia, histology, and probe path reconstruction

Mice were deeply anesthetized with a lethal dose of pentobarbital sodium/phenytoin before transcardial perfusion with phosphate-buffered 0.9% saline (PBS) followed by 4% paraformaldehyde (PFA). Brains were then extracted and stored in PFA at 4°C for 48 hours before transferring to PBS. Coronal sections of the brains around the probe recording sites were sliced in 70µm sections on a vibratome, mounted to coverslips, and counterstained with DAPI Fluoromount-G (SouthernBiotech, Alabama, USA).

Mounted sections were inspected under an epifluorescent microscope (Zeiss, AXIO Imager.Z1, California, USA) to resolve the presence of fluorescent dye. Slides were imaged at 5x magnification using DAPI, YFP, and Cy3 excitation/emission filter sets, and neighboring images were stitched together in Neurolucida Software (MBF Bioscience, Vermont, USA) to create one image per coronal section. Consecutive sections were mapped to the Allen Brain Reference Atlas common coordinate framework (Wang et al., 2020), and visible fluorescent probe tracks were manually trace (https://github.com/petersaj/AP_histology). A best-fit line through 3D reference atlas space was generated with the most ventral fluorescence signal defined as the probe tip. Electrode positions in atlas space were then assigned by projecting the probe electrode map (https://github.com/sotmasman/Silicon-microprobes) onto the best-fit line and the estimated depth.

### Spike sorting

Continuously sampled voltage data from the probe electrodes were spike sorted using Kilosort4 (Pachitariu et al., 2024) with default parameters. Rigid registration was used for drift correction. Following spike sorting, clusters were only included in further analyses if they had a signal-to-noise ratio greater than 5, less than 1% refractory period (1.5ms) violations, amplitude cutoff below 0.1, presence ratio higher than 0.5, and more than 500 total spikes. After applying these exclusion criteria, the remaining clusters were manually inspected in Phy, and clusters comprised of noise (e.g., non-physiological waveforms, near-equal amplitude across large channel span, abnormally stereotyped inter-spike-intervals) were excluded.

#### Electrical stimulation experiments

Bipolar stimulation electrodes were made by twisting stainless steel wire (A-M Systems, Washington, USA) and soldering them to separate pins (Digikey, Minnesota, USA). These electrodes were coated with DiI and lowered 0.8mm into S1 or V1 cortex bilaterally under isoflurane anesthesia (as in *Head-fixation and surface EEG electrode implant* in STAR Methods for surgical preparation details). Stimulation was performed with mice head-fixed on the polystyrene ball. During these sessions, stimulation was triggered every 20 seconds for 1 second at varying current intensities (0, 10, 30, 50, or 100 µA) at either 6 or 100Hz (pulse width: 1ms) using a stimulus isolator (WPI, Florida, USA). Mice underwent three recording sessions conducted on separate days. SWD start and end were labeled manually *post hoc*. Mice with fewer than five valid trials per condition (e.g., 0µA current at 6Hz is one condition) were excluded from the analysis. Correct electrode placement was verified by *post hoc* histology as above using histology of brain slices or with tissue clearing and subsequent lightsheet imaging. Briefly, after post-perfusion fixation in 4% PFA, brains were incubated in SHIELD OFF Solution (LifeCanvas Technologies, Massachusetts, USA) at 4°C for 72 hours and then transferred to SHIELD ON Solution at 37°C for 24 hours. Tissue clearing was performed in a SmartBatch+ (LifeCanvas Technologies, Massachusetts, USA) device for 24 hours according to the manufacturer’s standard protocol for mouse brain clearing. Samples were then incubated in EasyIndex (LifeCanvas Technologies, Massachusetts, USA) solution at 37°C for 48 hours to achieve tissue transparency and imaged with a Zeiss Lightsheet 7.

### QUANTIFICATION AND STATISTICAL ANALYSIS

#### Statistics software

GraphPad Prism (GraphPad Software, California, USA) was used to perform all t-tests, ANOVAs, and linear regressions. MATLAB (MathWorks, Massachusetts, USA) was used to perform D’Agostino K^2^ tests for skewness, Spearman rank correlations on sequential activity, pairwise correlations on neuronal spike trains, autocorrelations on proportional population vectors, and Pearson correlations on spike train pairs. RStudio (Posit Software, Massachusetts, USA) was used for linear mixed effects modeling for phase-locking analysis and analysis of S1/V1 stimulation effects on SWDs. Bonferroni-corrected p-values were used for Spearman rank correlations and subsequent ANOVAs to control for multiple comparisons. Benjamini-Hochberg adjusted p-values were used when classifying phase-locked neurons. When comparing phase-locking metrics (vector length and angle) across regions, the Tukey Test was used to compare all region pairs. When comparing stimulation current intensities to the control condition (0µA), Dunnet’s Test was used. Error bars in figures indicate 95% confidence intervals unless otherwise stated.

#### Semi-automated SWD detection

Candidate SWD events in surface EEG traces were detected by first computing a spectrogram with 1 second windows and 75% window overlap. Power between 4-8 Hz was calculated for all windows and a threshold equal to the 95^th^ percentile of the resultant distribution was used to identify candidate SWD events. Events briefer than 500ms were excluded and events separated by less than 1 second were merged. Troughs during these events were automatically detected with the *findpeaks* MATLAB function using a magnitude threshold of 3 standard deviations from the overall EEG voltage distribution and a minimum time difference of 50ms. These candidate events and troughs were manually verified. The time of the first trough was defined as the SWD start time. The time of the last trough was considered the SWD end time (**Figure 1C**).

#### Spike-SWD phase-locking

The instantaneous phase of an SWD was estimated by assigning individual troughs of SWDs the phase value of 0/360°, corresponding to the beginning of a cycle (**Figure 4B**). Intervals between troughs were divided into one hundred phase bins between 0 and 360°. Spike counts per neuron were binned according to these divisions, and the spike-count-weighted vector was computed as follows:

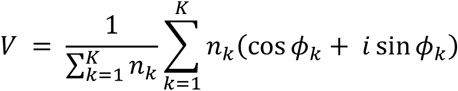

Therein, *nk* is the spike count in phase bin *k*, *φ_k_* is the phase angle of that bin, and *K* is the total number of phase angle bins. Vector (*V)* was normalized by the total spike count and phase angle (θ) and vector length (*ρ*) were taken as:

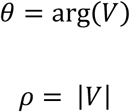

Therefore, the phase angle (θ) corresponds to point in the SWD cycle wherein a neuron fires, and the length (*ρ*) reflects the concentration of firing at this phase. Because of the normalization procedure, *V* was bounded between 0 and 1.

Linear mixed effects modeling (LME) was used to determine the effect of brain structure on phase-locking, as measured by mean vector lengths (**Figure 4E,K**). The following LME model was fit to our data in R using the ‘lmr’ function in the ‘lme4’ library:

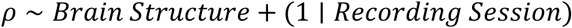

Here, *ρ* was the response variable, *Brain Structure* was included as a categorical fixed effect, and *Recording Session* was a random effect. Subsequent pairwise comparisons between regions were performed using the estimated marginal means of *ρ* from each structure, accounting for recording session variability. Tukey-adjusted p-values corrected for multiple comparisons (Table 1).

Significantly phased-locked neurons were determined by comparing observed vector lengths to those of null distributions generated by a random shifting procedure (**Figure S4**). For each neuron, spikes within a single SWD cycle were shifted by a random phase angle between 0° and 360° (**Figure S4A**). The random phase angle shift was generated independently for each cycle, and this shifting procedure was repeated 100,000 for each neuron (**Figure S4B**). The 100,000 resultant mean vector lengths then served as a null distribution (**Figure S4C**). The probability (*p*-value) of the observed vector length occurring within the null distribution was then calculated, and if it was less than a conservative specified significance level (*α* = 10^−4^), the neuron was considered significantly phase-locked (**Figure S1D**). Benjamini-Hochberg-adjusted *p*-values were used to correct for multiple tests in each brain. Overall, the motivation behind this shifting procedure was to preserve inter-spike intervals within SWD cycles but destroy cycle-to-cycle relationships thereby estimating the probability of getting the observed vector length by chance for each neuron.

#### Classification of spike waveforms

Spike waveform shapes were classified into two categories, putative excitatory and putative inhibitory, based on two features: trough-to-peak time and the half-amplitude duration (**Figure 4G**). Trough-to-peak times were measured by first finding the minimum voltage value (trough) of a neuron’s mean spike waveform recorded from the electrode with the highest amplitude. Then, the subsequent maximum (peak) was identified. The duration between these two values was the trough-to-peak time (**Figure 4G**). The half-amplitude was the duration between the time at which the mean spike waveform first passed half the value of its trough and then returned to that value after the trough (**Figure 4G**). Combining all waveforms from all recording sessions, two isolable clusters are present in this two-dimensional space (**Figure 4H**). The resultant two-dimensional distribution (trough-to-peak vs. half-amplitude duration) was then fit with a mixture of two Gaussians using *fitgmdist* and then separated into two clusters using cluster *in* MATLAB. The final decision boundary for waveform clustering can be seen in **Figure 4H**.

#### Sequential activity in SWD trough-centered (PETHs)

Peri-event time histograms (PETHs) were generated by binning spikes within a 100ms window around SWD troughs with a 1ms bin duration. PETHs were averaged across all troughs to generate a mean PETH for each neuron, temporally smoothed with a Gaussian window of 10ms, and then normalized to their own respective maximum value resulting in PETHs that could vary from 0 to 1. The center-of-mass time for each PETH was then defined:

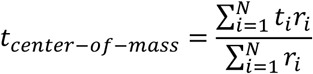

where *t_i_* is the time of bin *i*, *r_i_* is the normalized spike count in bin *i*, and *N* is the total number of bins (see **Figure 5B** for examples). Centers-of-mass were calculated for the PETHs from the first and second half of all SWD troughs in a recording session separately. Spearman rank correlations of these center-of-mass times were used then used to determine if sequential activity was preserved from the first to second half of troughs. If Spearman rank correlations were significant, sequential activity was considered preserved. Bonferroni-corrections were applied according to the number of tests in each brain region (e.g., if 10 tests were performed in region A, *α* becomes 0.05/10 or 0.005 for that region). For this analysis, only significantly phase-locked neurons were included. Only neurons recorded on the same shank were included in each observation because activity around SWD troughs may travel as a wave starting in different locations, thus disrupting inter-shank sequences (Luczak et al., 2007). Each shank in a recording session was considered an independent observation for this analysis (i.e., one session could yield two observations). Finally, only shanks with five or more neurons were included in this analysis.

#### Calculation of firing rates in different behavioral states

Each recording session was divided into 100ms time bins. Speed was calculated by summing the number of TTL signals during each bin, dividing by bin duration, and multiplying by distance per tick (0.957 cm) to return speed in cm/s units. Speed-by-time vectors were smoothed by convolution with a Gaussian kernel (500ms) to attenuate error introduced by quantization of the TTL signal from the rotary encoder. One of three labels was applied to each time point: “SWD”, “Immobile”, “Mobile”. All times during SWDs were labeled “SWD”. Periods containing no SWDs, during which there was also no movement for at least one second were labeled “Immobile”. All remaining times were labeled “Mobile”. Normalized firing rates over time (**Figures 2A** and **S1A**) were computed as in McCafferty et al. (2023). Thus, for each neuron, the mean peri-SWD firing rate vector was divided by its maximum value to normalize and constrain its range between 0 and 1. Initially, bin durations of 100ms (**Figure 2A**) were used, but durations of 500ms (**Figure S1A**) were also applied to reproduce the approach used by McCafferty (2023). Overall neuronal firing rates and firing rates in each behavioral state (**Figures 2G-I** and **S1B**) were calculated by dividing the total spike count in each behavioral state by the total duration of that state. Firing rates were statistically compared using paired t-tests when possible (**Figure S1B**) and one-way repeated measures ANOVAs with behavioral state as a factor and Bonferroni-corrected *post hoc* tests where appropriate (**Figures 2G-I** and **S1C**). In addition, to test whether SWDs appear during already ongoing immobility (**Figure 2F**), we used random temporal shifting to estimate the likelihood of finding our observed probability by chance. To this end, we circularly shifted the relative timing of SWDs a random amount between 30 and 600 seconds with respect to locomotion speed 10,000 times and computed a “excess probability” metric, which is the difference between the 95^th^ percentile value of the time-shifted null distribution and the actual observed probability (**Figure 2F**).

#### Pearson correlations of neuronal spike trains

Pearson correlations of neuronal spike trains were computed using MATLAB’s *corrcoef* function on pairs of spike trains during each Baseline and SWD period:

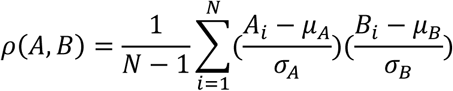

where *Ai* and *Bj* are the values of spike trains *A* and *B* at observation *i*, *N* is the total number of observations, *μ_A_* and *μ_B_* are the means of *A* and *B*, and *σ_A_* and *σ_B_* are their corresponding standard deviations. Pearson correlation coefficients (*p(A,B)*) were averaged across SWD and Baseline periods, excluding instances wherein one or both neurons did not fire. Finally, overall averages across all unique cell pairs within a given recording session were computed as follows:

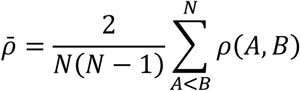

where *N* is the total number of neurons recorded. ρ̄ was computed separately for baseline and SWD periods and compared across conditions using paired t-tests (**Figure 3B**). Only sessions with at least five neurons in a brain area were included. The effect size (i.e., difference between mean correlations during baseline and SWD) was also compared using different bin durations when generating spike trains (**Figure S2A**).

#### Detection of HSEs

To quantify changes in the proportion of active neurons in the recorded population, we first discretized neuronal spike trains into 25ms bins in 5ms timesteps and found the proportion of neurons with at least one spike per bin across an entire recording session (***P****active*, Figure 6C, bottom):

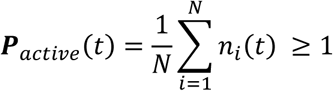

Therein, *ni(t)* is the spike count of neuron *i* in the current time bin *(t)*. Next, a distribution of ***P****active* during baseline periods was generated (**Figure 3C**, top left). Finally, a threshold equal to the 95^th^ percentile of the baseline ***P****active* distribution was used to find peaks separated by at least 25ms. These peaks were identified as high synchrony events (HSEs), and their rate of occurrence was compared across conditions using paired t-tests for each brain region (**Figure 3D**). Only sessions with at least five neurons in a brain area region included. The effect size (i.e., difference between the HSE rates during baseline and SWD) was also compared using different bin durations when generating ***P****active* (**Figure S2B**).

#### Quantifying differences between Pactive distributions

Because ***P****active* often showed profound silence, we also quantified the overall difference in ***P****active* distributions between baseline and SWD with no directionality bias using an approximation of the Wasserstein metric (Panaretos & Zemel, 2019). It was computed with the following equation:

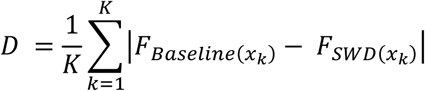

where *FBaseline* and *FSWD* are the cumulative distribution functions of ***P****active* in baseline and SWD periods, respectively, evaluated at *x*. Because this metric is the average absolute difference between the two distributions, we consider it a mean-normalized approximation of the 1D Wasserstein distance, which measures the minimal transport cost to convert one probability distribution into another (Panaretos & Zemel, 2019). *D* was calculated for each recording session, and *D* values were compared using one-way ANOVA with brain region as a factor and Bonferroni-corrected *post hoc* tests for all possible pairwise comparisons among brain regions. Only sessions with at least five neurons in a brain region were included.

#### Quantifying rhythmicity of population activity

The rhythmicity of ***P****active* was quantified using an oscillation index (OI, Kleiman-Weiner et al., 2009). First, the autocorrelation of ***P****active* during baseline and SWD periods was computed with the *xcorr* MATLAB function with normalization which implements the following operation:

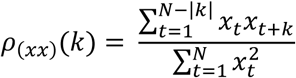

where *p(xx)k* is the normalized autocorrelation at lag *k*, *x* is the value of ***P****active* at time *t*, and *N* is the total number of time points. This was performed for every baseline and SWD period and a weighted average was taken across all corresponding periods (i.e., weighted by the duration of the period). After finding *p(xx)k*, OI was calculated by finding the first peak at *x > 0* and subtracting from that the trough between it and the central peak at *x = 0* (**Figure 3F**). Therefore, the final OI value is bounded between 0 (no periodicity) and 1 (perfect periodicity). OI was compared across conditions using separate paired t-tests for each brain region (**Figure 3G**).

#### Statistical analysis for electrical stimulation experiments

To assess the effect of stimulation current intensity on SWD induction and termination probabilities, we fit an LME model with trial outcome as the dependent variable, stimulation as a fixed effect, and mouse identity as a random intercept to account for multiple sessions from each mouse:

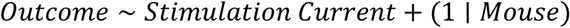

Multiple comparisons to the 0µA baseline value were performed with Dunnett’s *post hoc* tests.

#### Multi-unit activity (MUA) for laminar activation analysis

MUA was computed by first downsampling the raw 30kHz data to 10kHz. Then, dead and noisy electrodes (i.e., those with too little or too much voltage fluctuation) were removed by finding the standard deviation of voltages over time for each electrode and excluding electrodes that fell outside of the 5^th^ to 95^th^ percentile range. Remaining channels from each shank were referenced to a within-shank common average to remove any extant common-mode noise. Signals were then high-passed filter at 300Hz and a signal median-based detection threshold (Quiroga et al., 2004) was applied to the rectified signal. The detection threshold is given by:

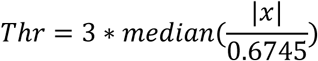

where *x* is the high-pass filtered signal. For this analysis, single shanks were considered independent observations and only shanks that had electrodes spanning all cortical layers in S1 or V1 were included. Center-of-mass times were derived from MUA SWD trough-centered PETHs averaged within layers (see *Sequential activity in SWD trough-centered (PETHs)* in STAR Methods for explanation of PETH calculations). Center-of-mass times across layers were analyzed with one-way repeated measures ANOVA and Bonferroni *post hoc* comparisons to compare times from individual layers to each other.

## KEY RESOURCES TABLE

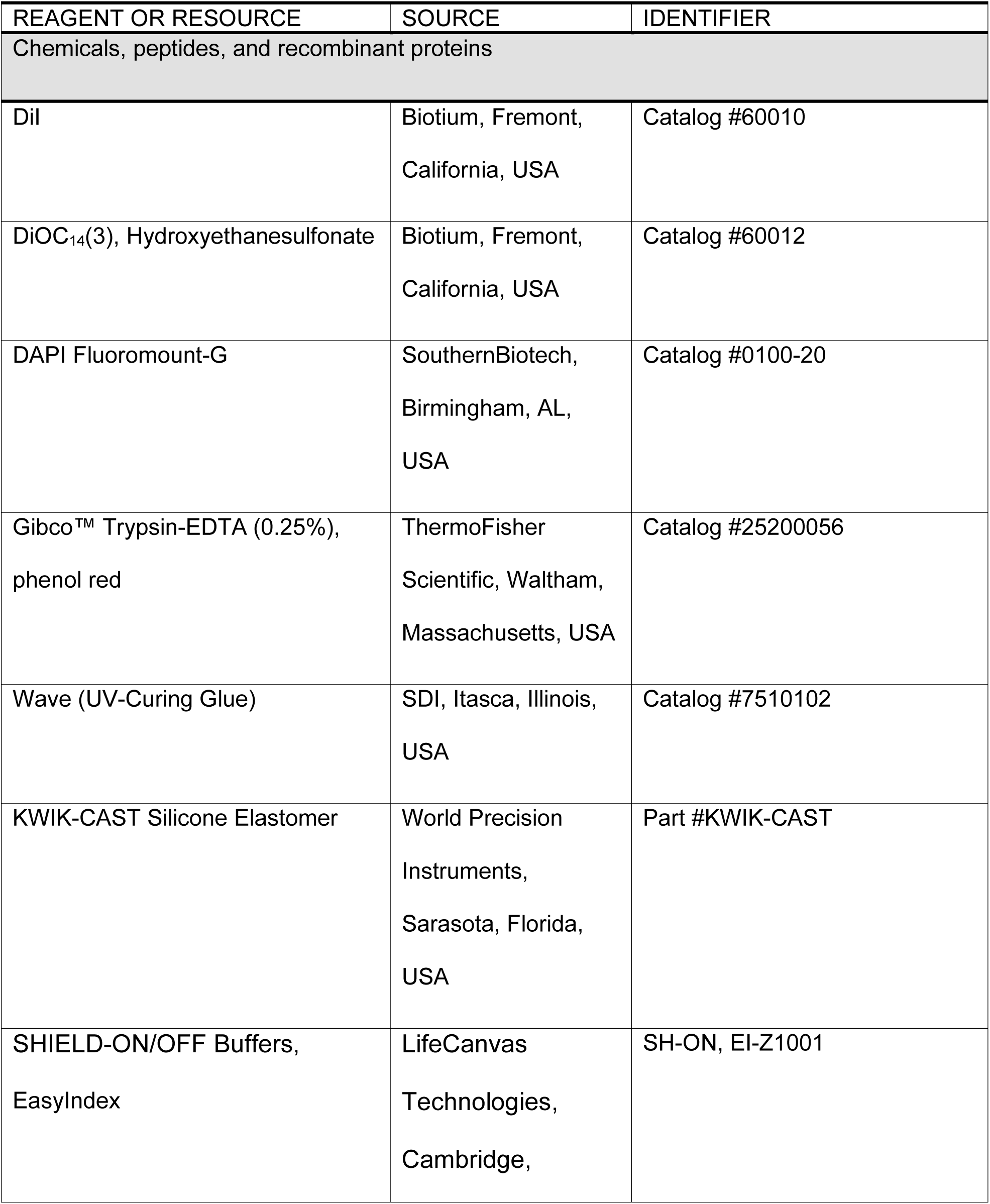
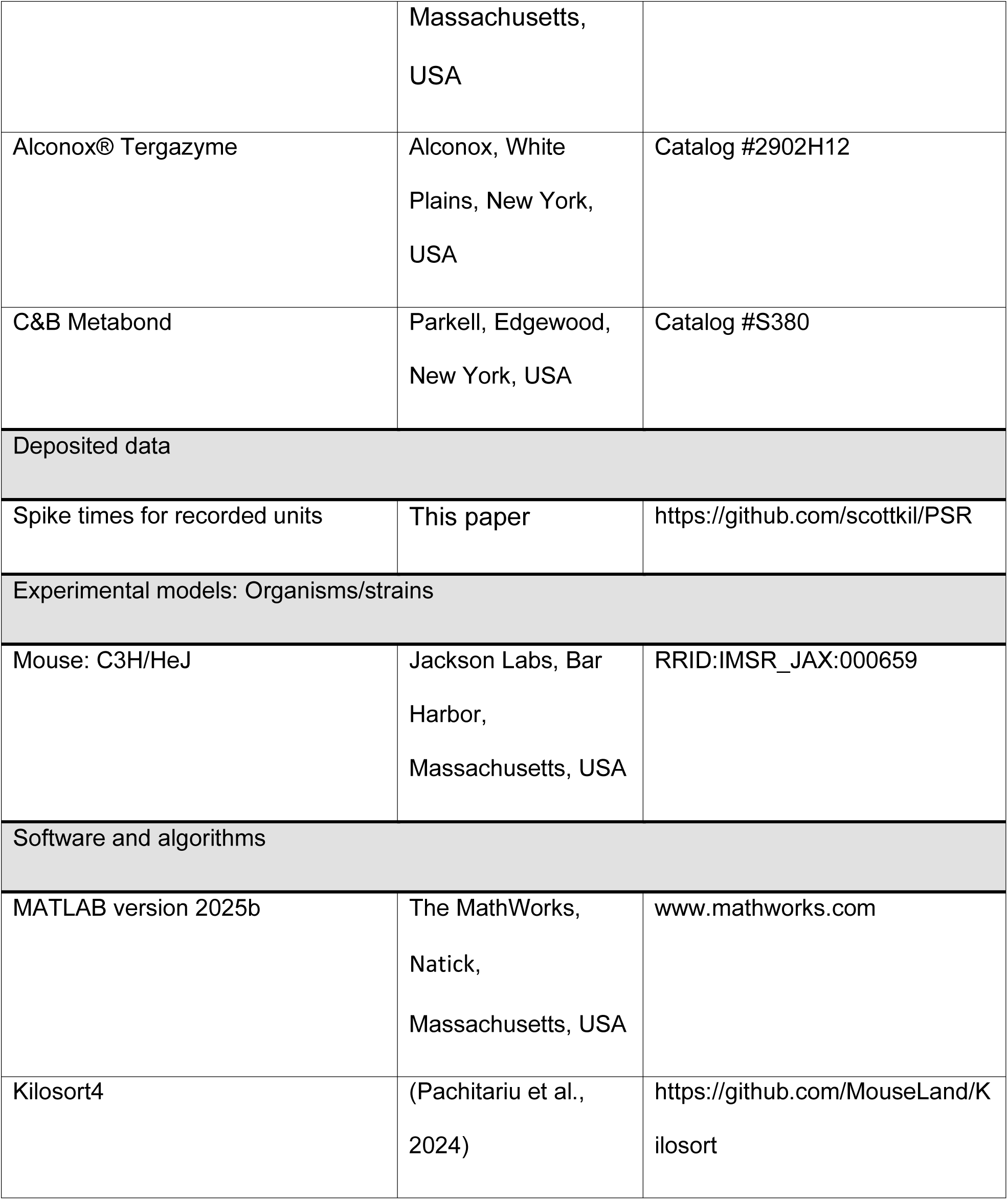
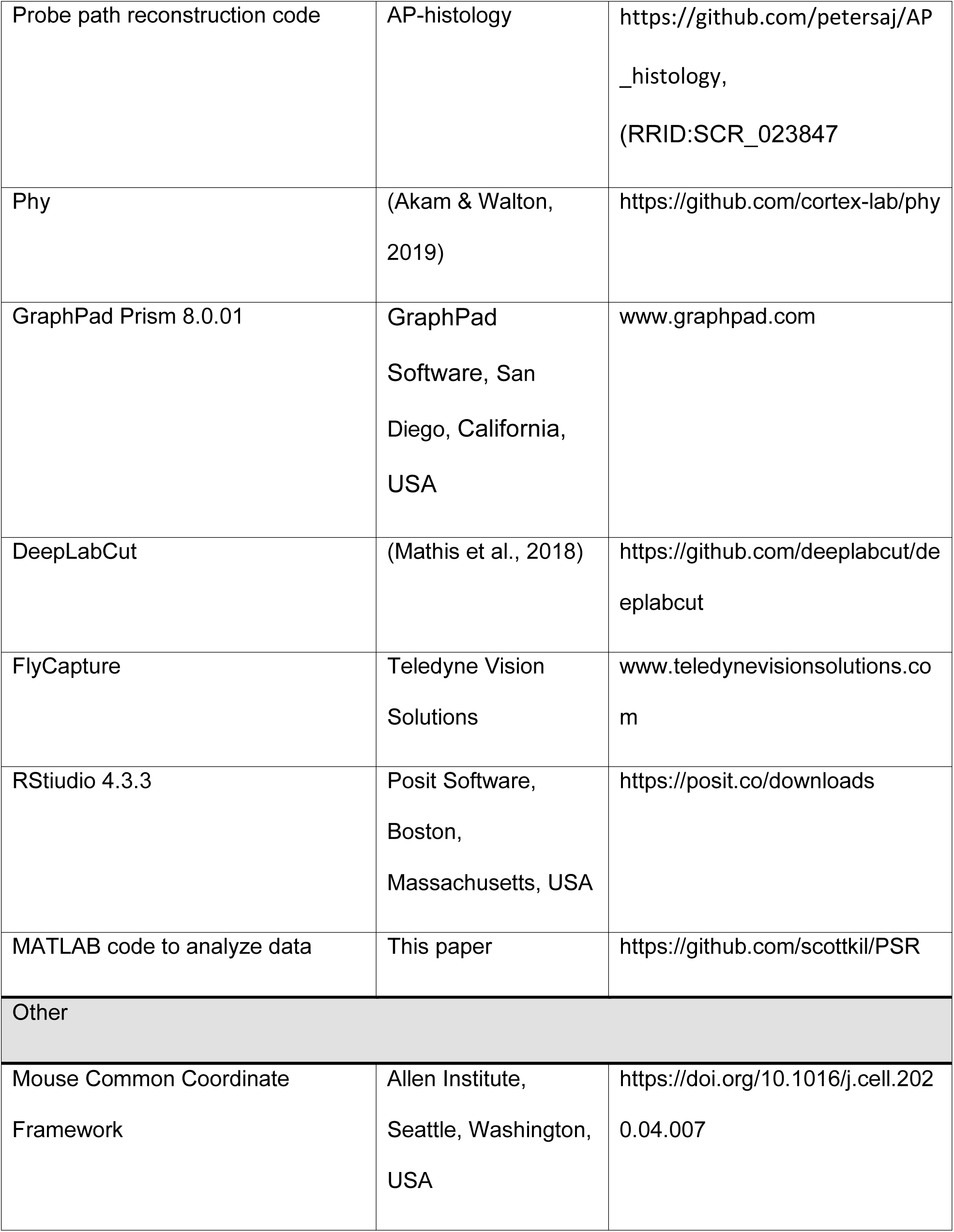
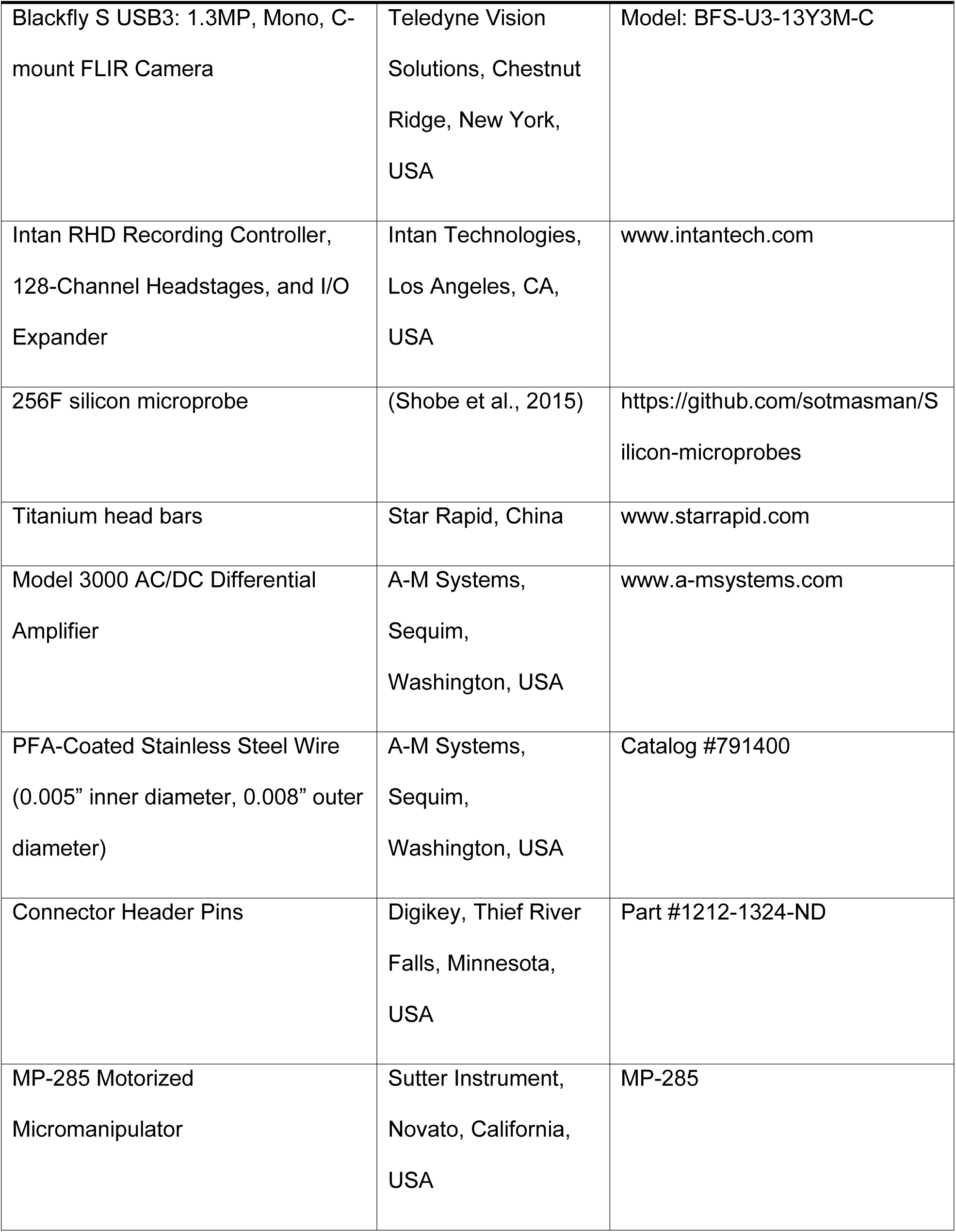
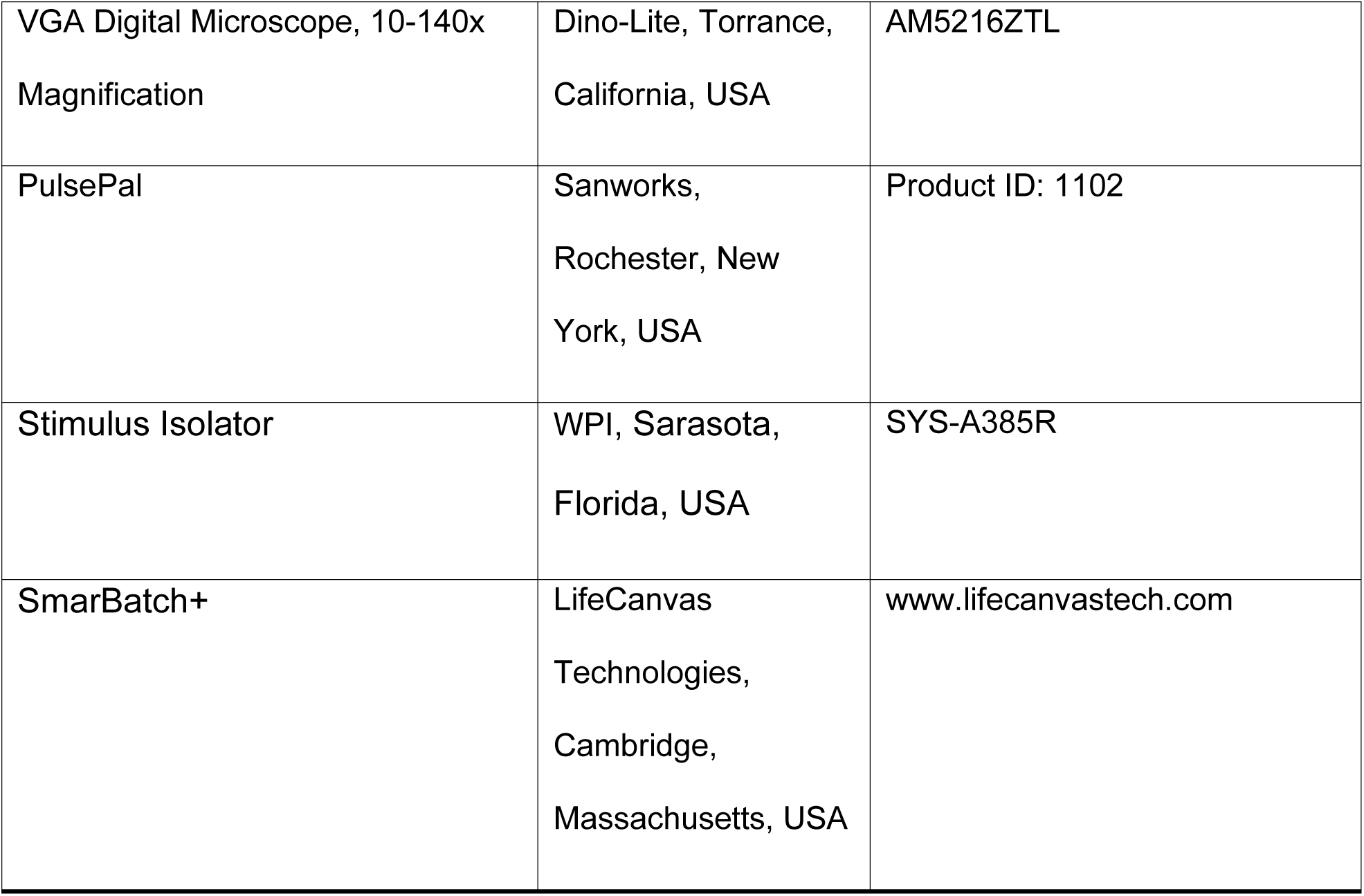

