## Supplementary Material for "Spike-wave discharges reflect widespread, but non-uniform, cortical hypersynchrony during immobility"

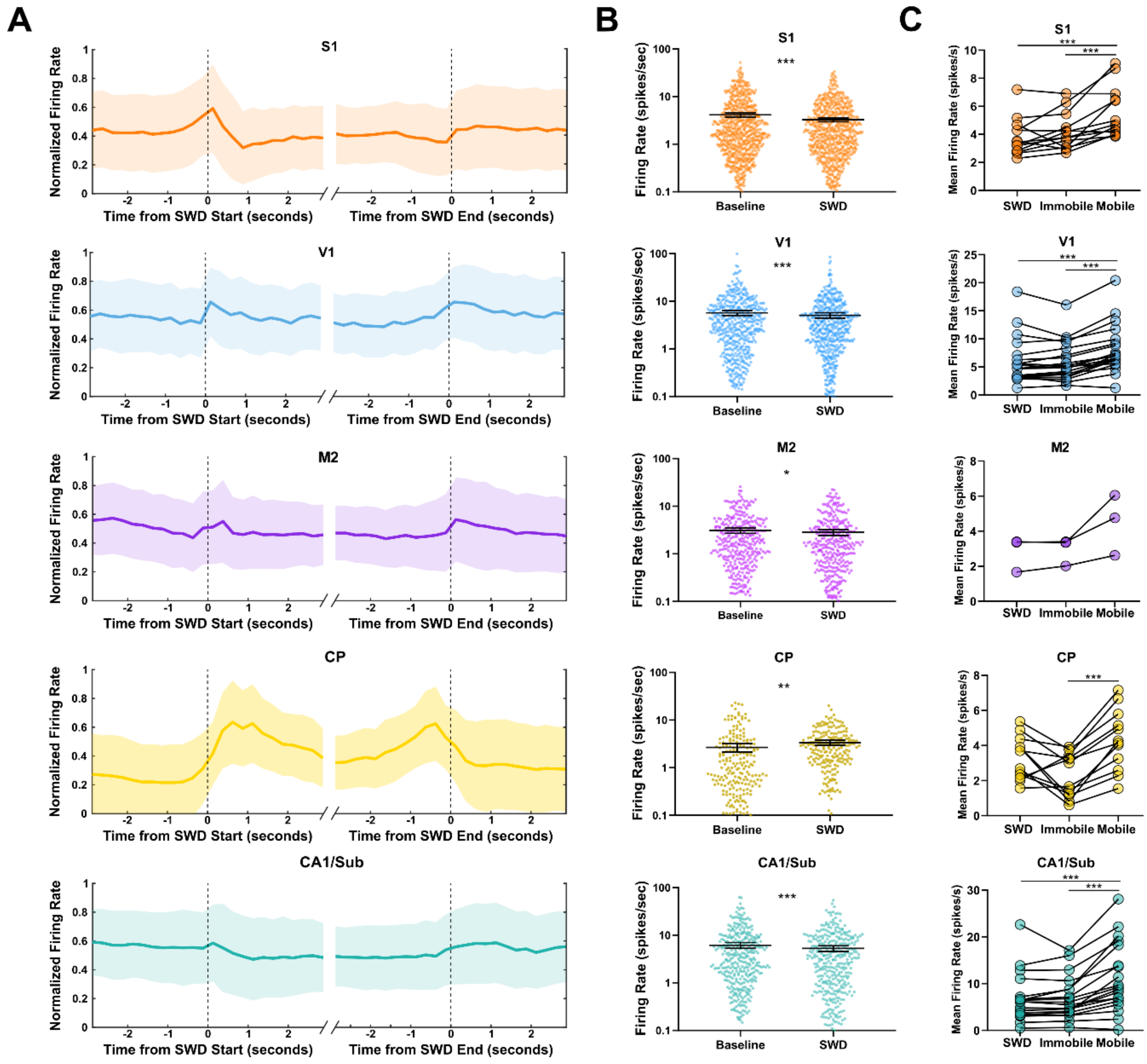


**Figure S1. Firing rates with longer bin durations, overall Baseline and SWD rates, and session means. (A)** Average normalized firing rates for all recorded structures relative to the starts (left) and ends (right) of SWDs using 500ms bins. Shading is ± SD. **(B)** Firing rates during SWDs and all non-ictal, baseline periods (combined Immobile and Mobile). Horizontal black lines and bars show mean and 95% CI. **(C)** Average firing rates per session in different behavioral states. * $p < 0.05$, ** $p < 0.01$, *** $p < 10^{-3}$.

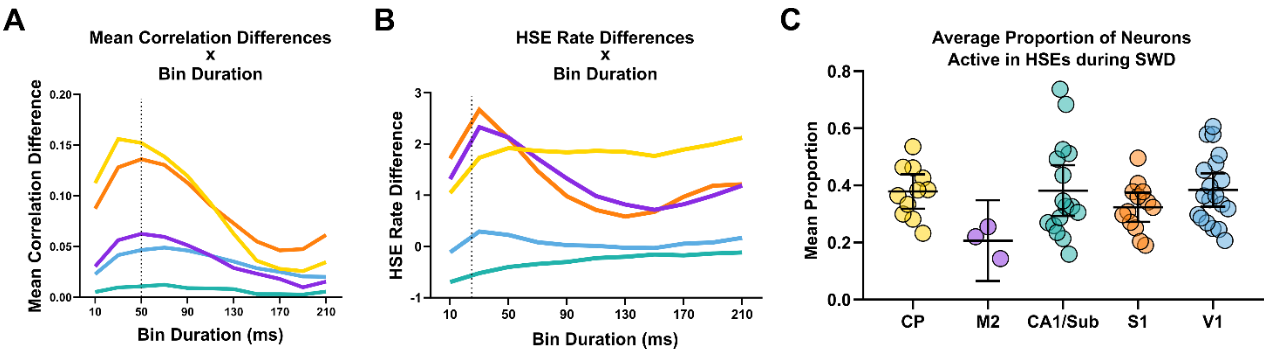


**Figure S2. Spike train correlations and HSE rate with different $P_{active}$ bin durations. (A)** Differences between pairwise spike train correlations from baseline and SWD using increasing bin durations when computing $P_{active}$. Vertical black dotted line indicates the bin duration used in **Figure 2A,B**. **(B)** Differences between HSE rates during baseline and SWD using increasing bin durations when computing $P_{active}$. Vertical black dotted line indicates the bin duration used in **Figure 2C,D**, and **E**. **(C)** Average proportion of neurons active in HSEs occurring during SWDs. Error bars indicate 95% confidence intervals.

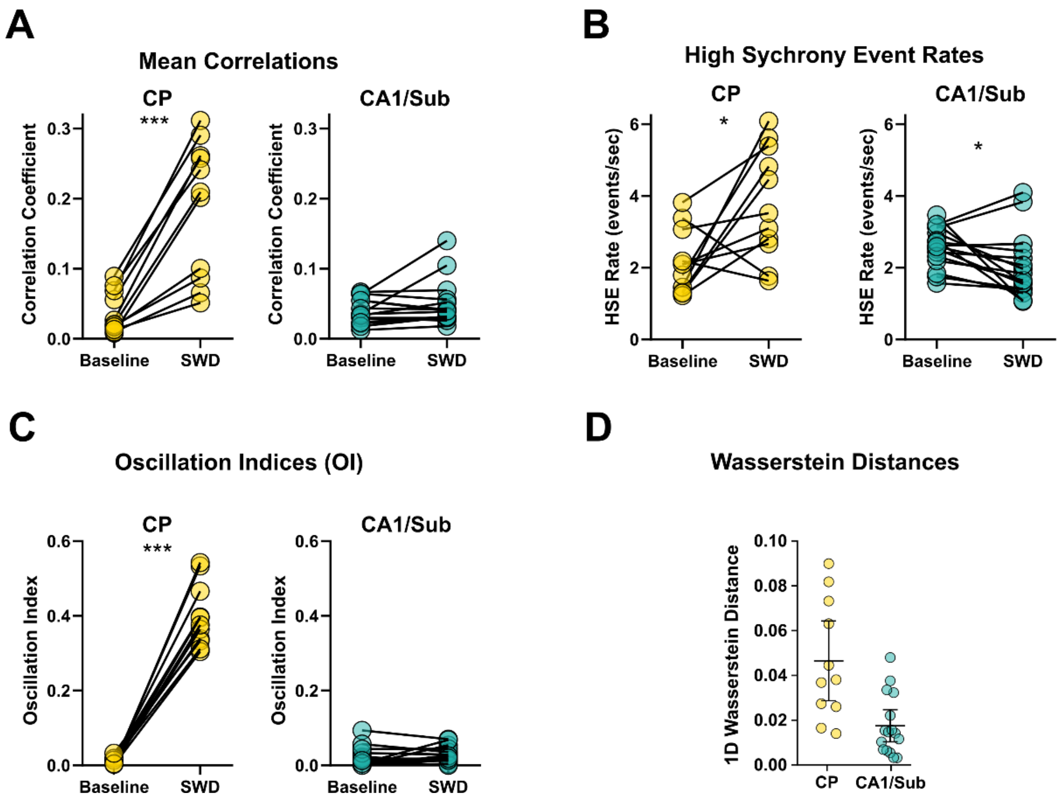


**Figure S3. Synchrony metrics for CP and CA1/Sub. (A)** Mean correlation coefficients during baseline and SWD. Each session is represented by a pair of connected circles. **(B)** HSE rates during baseline and SWD. **(C)** OI during baseline and SWD. **(D)** 1D Wasserstein distances between $P_{active}$ distributions. Each circle represents one recording session. * $p < 0.05$, ** $p < 0.01$, *** $p < 10^{-3}$. CP: n=11; CA1/Sub: n=16.

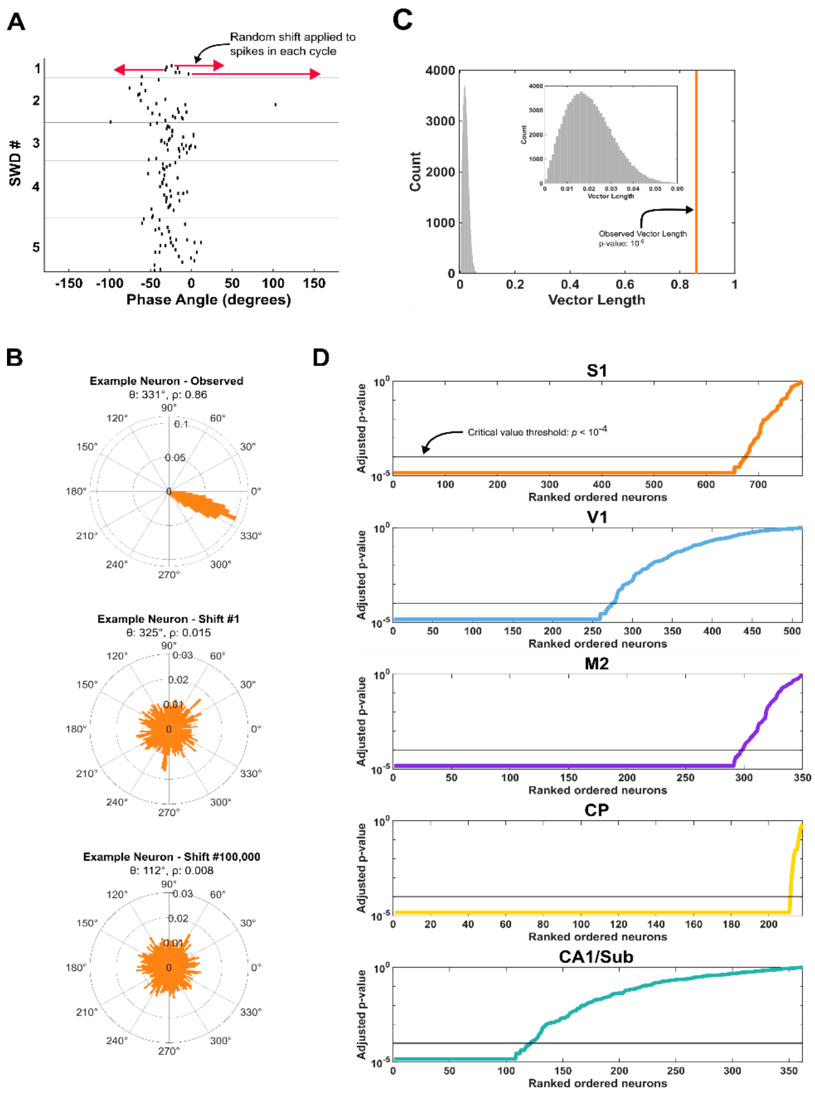


**Figure S4. Determination of significant phase-locking for single neurons. (A)** Spike-phase raster plot for one significantly phase-locked neuron during five SWDs. The y-axis corresponds to individual SWD cycles during each of five distinct SWDs separated by horizontal lines. X-axis corresponds to the phase angle within a given SWD cycle. Horizontal red arrows indicate the random shifting procedure which were applied to groups of spikes within cycles separately. Only three random shifts are shown here for simplicity, but different random time shifts were applied to all cycles. This neuron showed obvious phase locking in the non-shifted spike times (i.e. black raster lines). **(B)** Polar histograms generated from the actual spike phase angles (top) and randomly shifted spike phase angles (bottom two). θ and ρ indicate the mean vector angle and length, respectively. Note the vastly different radial scales in the top histogram relative to the bottom two histograms due to the destruction of phase-locking by the random shifting procedure. Also note the relatively uniform distribution of phase angles in the bottom two histograms relative to the highly constrained angles of the top histogram. **(C)** Histogram of the vector lengths produced by the 100,000 random phase angle shifts. The orange vertical line corresponds to the observed vector length for this neuron and was larger than all of the shifted vector lengths. The resultant probability of finding observed vector length is therefore $10^{-5}$. **(D)** Rank-ordered, Benjamini-Hoch adjusted p-values for all neurons plotted separately for each brain region. Horizontal lines indicate the p-value threshold used to determine if a neuron was significantly phase-locked. Neurons with p-values less than $10^{-4}$ were beyond this threshold. Note the logarithmic y-axis in these plots

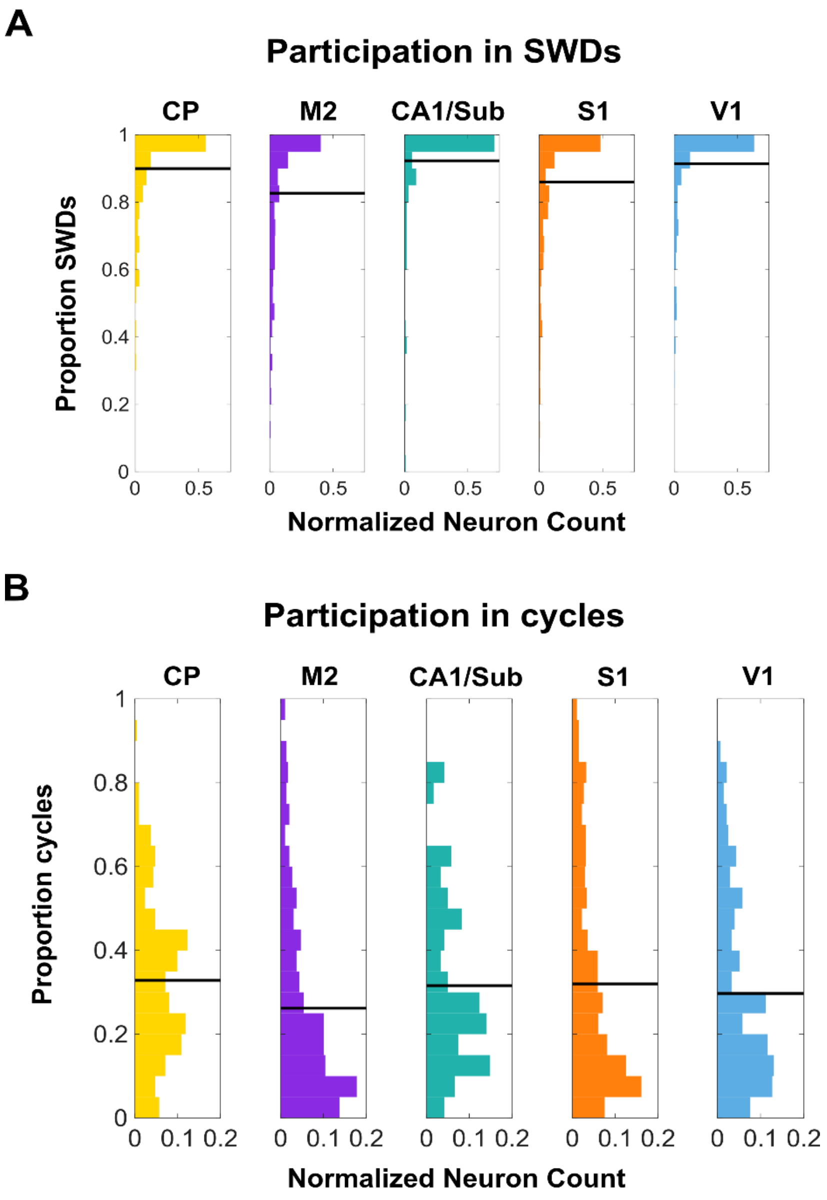


**Figure S5. Neuronal participation during entire SWDs and during individual SWD cycles. (A)** Histograms of neuronal participation during at least some portion of an entire SWD. Neurons were considered to be participating during an SWD if they fired at least once during the SWD. Population means are represented by horizontal black lines. Only significantly phase-locked neurons were included in this analysis. **(B)** Histograms of neuronal participation during individual SWD cycles. Neurons were considered to be participating during a cycle if they fired within a 60ms window around SWD troughs (±30ms). Means are represented by horizontal black lines. Only significantly phase-locked neurons were included in this analysis.

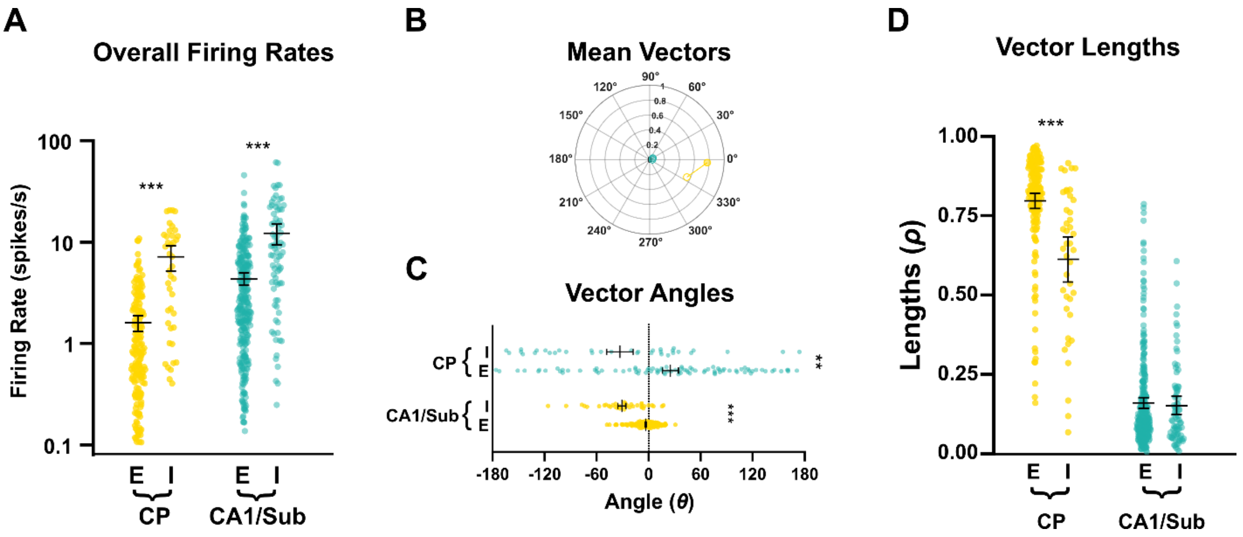


**Figure S6. Firing rates and phase-locking for inhibitory and excitatory neurons in CP and CA1/Sub.** **A)** Overall firing rates of putative excitatory and inhibitory neurons. **B)** Mean spike-SWD vector separated by putative neuron type. Open circles represent mean vectors for putative inhibitory neurons while shaded circles are those of excitatory neurons. Putative inhibitory neurons have earlier mean vector angles. **C)** Vector angles of putative excitatory and inhibitory neurons. Bars indicate the 95% confidence interval. **D)** Vector lengths of putative excitatory and inhibitory neurons.

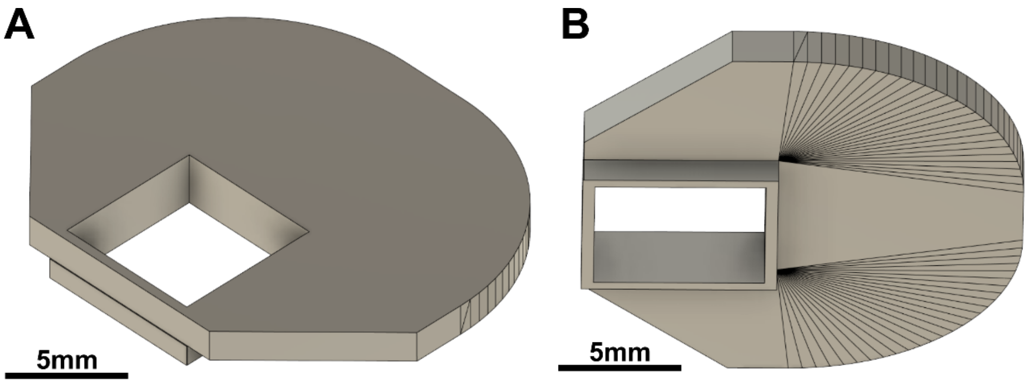


**Figure S7. 3D rendering of protective chamber used for silicon probe recordings. A)** Top of the chamber viewed from a posterolateral angle. **B)** Bottom of the chamber viewed from a lateral angle. See *Materials availability* in STAR Methods for design files.

**Supplementary Table S1. Mouse and SWD statistics by cortical target**

| Cortical Target | # Recording Sessions | Mouse Age (days) | SWD Rate (Hz) | SWD Duration (s) | SWD Peak Frequency (Hz) |
|---|---|---|---|---|---|
| **M2** | 3 | 78 ± 1.0 | 1.3 ± 0.4 | 3.8 ± 0.83 | 5.9 ± 0.3 |
| **S1** | 13 | 62.1 ± 7.2 | 1.2 ± 0.7 | 3.4 ± 1.1 | 6.0 ± 0.6 |
| **V1** | 21 | 62.6 ± 5.1 | 1.1 ± 0.6 | 3.6 ± 0.9 | 6.0 ± 0.4 |

**Supplementary Table S2. Number of recording sessions and neurons by brain region and layer**

| | Primary Somatosensory (S1) | Primary Visual (V1) | Secondary Motor (M2) | Caudoputamen (CP) | CA1/Sub |
|---|---|---|---|---|---|
| # Recording Sessions | 13 | 21 | 3 | 12 | 19 |
| # Total Neurons | 785 | 513 | 350 | 218 | 362 |
| Layer II/III | 120 | 67 | 0 | | |
| Layer IV | 94 | 47 | | | |
| Layer V | 135 | 131 | 80 | | |
| Layer VI | 353 | 204 | 270 | | |
| Unassigned Layer | 83 | 64 | 0 | | |

**Supplementary Table S3. Participation rates per-SWD and per-cycle for significantly phase-locked neurons**

| Brain Region | # Significantly phase-locked neurons | Mean per-SWD participation rate | D'Agostino $K^2$ *Statistic* | Mean per-cycle participation rate | D'Agostino $K^2$ *Statistic* |
|---|---|---|---|---|---|
| **CP** | 211 | 90% | 87.2*** | 33% | 8.2* |
| **M2** | 298 | 83% | 60.1*** | 26% | 48.3*** |
| **CA1/Sub** | 121 | 92% | 112.9*** | 32% | 11.2** |
| **S1** | 675 | 86% | 219.0*** | 32% | 70.1*** |
| **V1** | 275 | 91% | 132.5*** | 30% | 26.1*** |

* $p < 0.05$, ** $p < 0.01$, *** $p < 0.001$
